# Pathological modeling of TBEV infection reveals differential innate immune responses in human neurons and astrocytes that correlate with their susceptibility to infection

**DOI:** 10.1101/819540

**Authors:** Mazigh Fares, Marielle Cochet-Bernoin, Gaëlle Gonzalez, Claudia N. Montero-Menei, Odile Blanchet, Alexandra Benchoua, Claire Boissart, Sylvie Lecollinet, Jennifer Richardson, Nadia Haddad, Muriel Coulpier

**Affiliations:** UMR1161 Virologie, Anses, INRA, Ecole Nationale Vétérinaire d’Alfort, Université Paris-Est, Maisons-Alfort, France; MRC-University of Glasgow Centre for Virus Research, Glasgow, Scotland, United Kingdom; CRCINA, UMR 1232, INSERM, Université de Nantes, Université d’Angers, Angers F-49933, France; CHU Angers, Centre de Ressources Biologiques, BB-0033-00038, Angers, France; CECS, I-STEM, AFM, Evry, France; UMR BIPAR 956, Anses, INRA, Ecole Nationale Vétérinaire d’Alfort, Université Paris-Est, Maisons-Alfort, France

**Author notes:** Author contribution* Conceptualization : MF and MC; Development or design of methodology : MF and MC; Validation : MF and MC; Formal analysis : MF; Investigation : MF, MCB, GG; Resources : CNMM, OB, AB, CB and SL; Original draft preparation : MF and MC; Writing, review and Editing : JR and NH; Visualization : MF and MC; Supervision : MC; Project administration : MC; Funding acquisition : NH and MC.

## Abstract

Tick-borne encephalitis virus (TBEV) is a member of the *Flaviviridae* family, *Flavivirus* genus, which includes several important human pathogens. It is responsible for neurological symptoms that may cause permanent disability or death, and, from a medical point of view, is the major arbovirus in Central/Northern Europe and North-eastern Asia. TBEV tropism is critical for neuropathogenesis, yet, little is known about the molecular mechanisms that govern the susceptibility of human brain cells to the virus. In this study, we sought to establish and characterize a new *in vitro* model of TBEV infection in the human brain and to decipher cell type-specific innate immunity and its relation to TBEV tropism and neuropathogenesis. We showed that infection of neuronal/glial cultures derived from human fetal neural progenitor cells (hNPCs) mimicked three major hallmarks of TBEV infection in the human brain, namely, preferential neuronal tropism, neuronal death and astrogliosis. We also showed that these cells had conserved their capacity to build an antiviral response against TBEV. TBEV-infected neuronal/glial cells, therefore, represented a highly relevant pathological model. By enriching the cultures in either human neurons or astrocytes, we further demonstrated qualitative and quantitative differential innate immune responses in the two cell types that correlated with their particular susceptibility to TBEV. Our results thus reveal that cell type-specific innate immunity is likely to contribute to shaping TBEV tropism for human brain cells. They offer a new *in vitro* model to further study TBEV-induced neuropathogenesis and improve our understanding of the mechanisms by which neurotropic viruses target and damage human brain cells.

**Author summary:** Tick-borne encephalitis virus (TBEV), a neurotropic *Flavivirus* that is responsible for encephalitis in humans, is of growing concern in Europe. Indeed, over the last two decades the number of reported cases has continuously increased and the virus has spread into new geographical areas. Whereas it is well established that neurons are the main target of TBEV in the human brain, the mechanisms that underlie this preferential tropism have not yet been elucidated. Here, we used neuronal/glial cells derived from human fetal neural progenitors to establish and characterize a new *in vitro* pathological model that mimics major hallmarks of TBEV infection *in vivo*; namely, neuronal tropism, neuronal death and astrogliosis. Using this highly relevant model, we showed that human neurons and astrocytes were both capable of developing an innate immune response against TBEV, but with dissimilar magnitudes that correlated with differential susceptibility to TBEV. Our results thus revealed that TBEV tropism for subsets of human brain cells is likely to depend on cell-type specific innate immunity. This improves our understanding of the mechanisms by which neurotropic viruses target and damage human brain cells and may help guide development of future therapies.

## Introduction

Tick-borne encephalitis virus (TBEV) belongs to the genus *Flavivirus* (family *Flaviviridae*), whose members include several important human pathogens transmitted by arthropods, such as Japanese encephalitis virus (JEV), West Nile virus (WNV), Zika virus (ZIKV) and Powassan virus (POWV). From a medical point of view, TBEV is the most important arbovirus in Europe and North-eastern Asia. Its endemic zone spreads from Northern, Central and Eastern Europe to Far East Asia (1). It induces a range of symptoms from mild flu-like symptoms to severe encephalitis and paralysis, often with long term neurological sequelae (2). The incidence of the disease has increased in recent decades and autochtonous cases are regularly reported in new areas of Western Europe, reflecting an expansion to non-endemic areas (3). Despite commercialization of an effective vaccine (4), between 8000 and 13000 annual cases of tick-borne encephalitis have been reported worldwide since the 1990s (5). No therapy is currently available (6).

TBEV is usually transmitted to humans from infected ticks, mainly of the *Ixodes* family, but may occasionally be acquired by consumption of unpasteurized dairy products from infected livestock (7–9). Upon inoculation into the human skin, initial infection and replication occurs in local dendritic cells (DCs). DCs are believed to transport the virus to draining lymph nodes from which it spreads into the bloodstream and induces viremia. It may then cross the blood brain barrier and cause widespread lesions in the brain. These include inflammatory changes, neuronal damage and glial reactivity in several brain areas, including the spinal cord, brainstem, cerebellum and striatum (10, 11). Neurons are the primary target of infection (12) but other cells in the CNS may contribute to TBEV-induced neuropathogenesis. Both infiltrating immunocompetent cells, mainly CD8^+^ T cells, and resident glial cells, such as astrocytes and microglial cells, have been shown to play a role (13, 14). Neuronal damage may thus be mediated directly by viral infection or indirectly by infiltrating immunocompetent cells, inflammatory cytokines and activated resident glial cells.

The innate immune response is the first line of defense against viral infection. Type I interferons (IFNs) are of particular importance in this process. Through binding to the IFN alpha/beta receptor (IFNAR), they act via autocrine or paracrine signaling (15–17) and trigger the activation of a large number of interferon-stimulated genes (ISGs) that can inhibit almost every step of viral life cycle (18). In recent years, it has become clear that parenchymal cells of the central nervous system (CNS) play a major role in the development of the innate immune response and the protection of infected individuals after CNS infection (19–24). Neurons and astrocytes are not passive targets, as they are known to produce and respond to type I IFNs. Nevertheless, the innate immune programs activated in these cell types during TBEV infection and their impact on viral tropism and neuropathogenesis remain poorly known.

Animal models have been widely used to elucidate the cellular and molecular mechanisms of TBEV-induced neuropathogenesis (2). Nevertheless, the results obtained from such studies may be difficult to transpose to human neuropathogenesis, as human anti-viral responses differ substantially from those of other mammalian species (25, 26). The biological relevance of models based on human CNS cells is thus increasingly recognized. These include neuronal/glial cell lines, primary neuronal/glial cells from human fetuses and, more recently, neuronal/glial cells derived from fetal neural progenitors (hNPCs), embryonic (hESC) or induced pluripotent stem cells (hiPSCs). While primary human CNS cells are physiologically more relevant than cell lines, their use is limited by the difficulty in gaining access to cell sources. On the other hand, neuronal/glial cultures derived from neural progenitors are not only physiologically relevant, but also have the advantage of being available on demand. In recent years, they have become important tools to study neurotropic viruses (27).

The goal of this study was, first, to set up and characterize a new *in vitro* model of TBEV infection using complex co-cultures of hNPC-derived neuronal/glial cells, and second, to decipher cell-specific anti-TBEV immunity in the human CNS and its relation to TBEV tropism and neuropathogenesis. We showed that *in vitro* TBEV infection mimics several hallmarks of *in vivo* infection, including marked neuronal tropism and neuronal death, limited astrocyte susceptibility, astrogliosis and induction of an antiviral response. Moreover, we demonstrated differential qualitative and quantitative antiviral capacities in human neurons and astrocytes that are correlated with their susceptibility to TBEV infection. Finally, we showed that human astrocytes exert a protective effect on neighboring TBEV-infected neurons.

## Results

### TBEV infects brain cells differentiated from human fetal neural progenitors

HNPCs and their derived neuronal/glial cells were previously set up in our laboratory for the study of Borna disease virus, a neurotropic virus that belongs to the *Bornaviridae* family (28, 29). Here, we used the same hNPCs prepared according to the experimental steps summarized in Fig. 1A. HNPCs were differentiated for 13 days, by which time all neurons were generated (29), before infection with the TBEV-Hypr strain at MOI 10^-2^. Infected neuronal/glial co-cultures were then analyzed over time. We first examined the capacity of the virus to infect, replicate and disseminate within the co-culture. Examination of cells immunostained with an antibody specific for domain 3 of the TBEV envelope protein (TBEV-E3), at 24 and 72 hours post-infection (hpi), revealed that TBEV entered human brain cells and spread within the co-culture (Fig. 1B). Enumeration of infected cells from 14 hpi to 7 dpi showed that 7.3±0.7% of cells were indeed infected at 14 hpi whereas 45.0±4.0% were infected at 72 hpi, at the peak of infection (Fig. 1C). At a later time, 7 dpi, the number of infected cells decreased. A similar pattern of infection was observed when the viral RNA was quantified by RT-qPCR, whether in the supernatant or intracellularly, from 14 hpi to 7 dpi (Fig. 1D) with an increase in viral RNA observed up to 48-72 hpi followed by a decrease from 96 hpi to 7 dpi. This confirmed active replication of the virus in hNPC-derived brain cells. Quantification of viral titer by end-point dilution further showed that infectious particles were released into the supernatant at 48 and 72 hpi (Fig. 1E). Thus, the infection, replication and dissemination of virus were efficient in hNPC-derived neuronal/glial cells.

**Figure 1.**
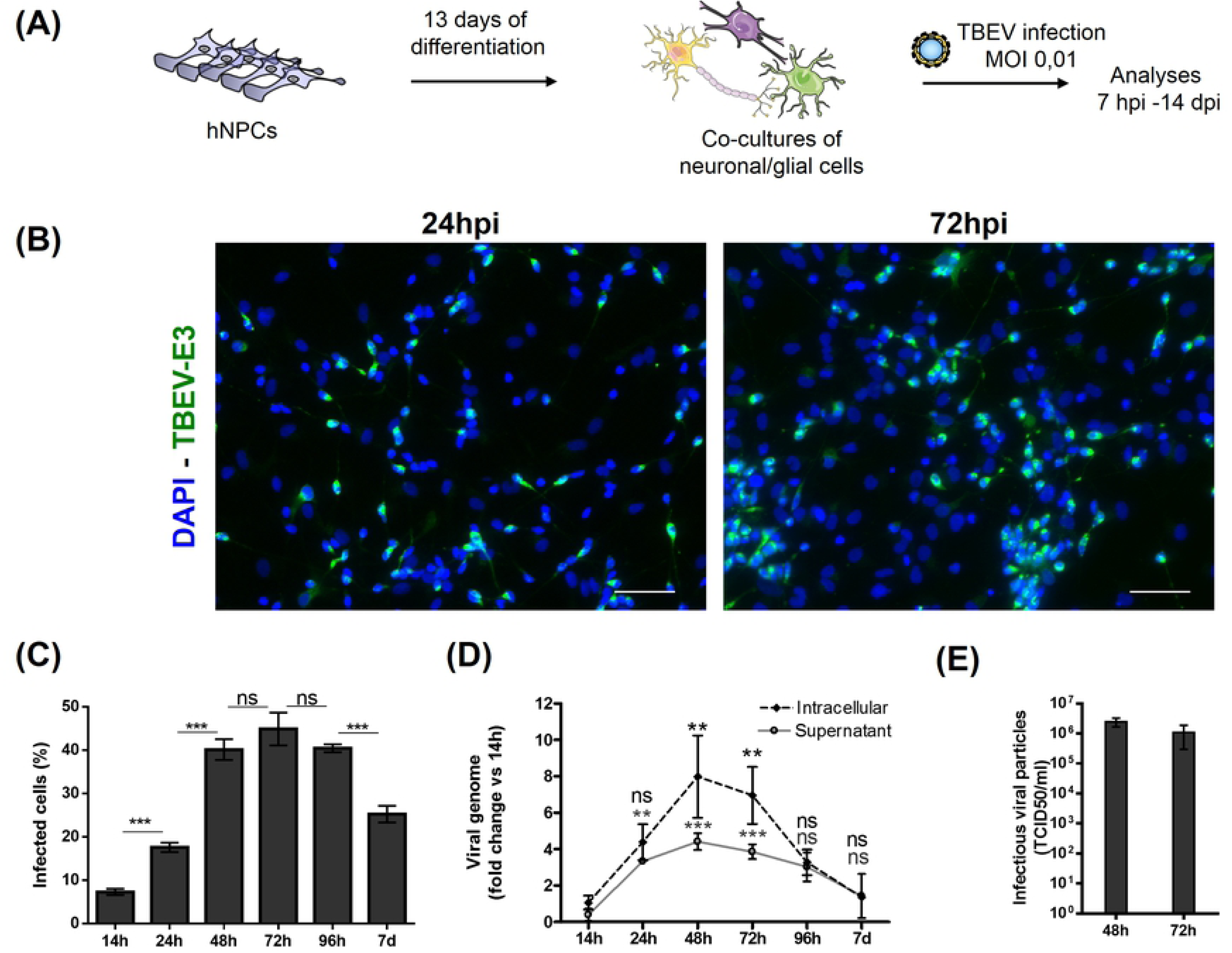
TBEV infects, replicates and spreads in hNPC-derived brain cells. (A) Schematic representation of the experimental procedure. (B) Immunofluorescence labeling of differentiated hNPCs 24 h and 72 h following TBEV infection. An antibody directed against the domain 3 of the viral envelope (TBEV-E3, green) revealed infected cells. Nuclei were stained with DAPI (blue). (C) Enumeration of infected cells based on immunofluorescence labeling using an ArrayScan Cellomics instrument. (D) RNA from the supernatant and cell lysate of infected cells was analyzed by RT-qPCR to determine viral replication. (E) Supernatant was collected at the peak of infection and titrated by end-point dilution (TCID50) on VERO cells. Results are representative of 3 independent experiments performed in triplicate. Data are expressed as the mean ± SD. Statistical analysis was performed using one-way ANOVA (Bonferroni’s Multiple Comparison Test) with Graphpad Prism V6.0.1, ns=non-significant (p>0.05); **=p<0.01, ***=p<0.001. Scale bar=100µm.

### TBEV infects human neurons, astrocytes, and oligodendrocytes

We next sought to determine which neural subsets were infected by TBEV in neuronal/glial co-cultures. We had previously shown that, upon growth factor withdrawal, hNPCs differentiated into neurons and astrocytes (28, 29). Oligodendrocytes, the third cell type that can be generated by differentiation of hNPCs, were not taken into consideration. To gain in precision, in the present study we enumerated all 3 cell types, based on immunofluorescence staining (S1A Fig), 13 and 21 days after the onset of differentiation. Automatic enumeration of immunostained cells with antibodies directed against HuC/HuD (nuclear markers for neurons) and OLIG2 (nuclear marker for oligodendrocytes) revealed a population composed of 77.0±3.2% (d13) and 74.1±5.4% (d21) neurons and 1.4±1.0% (d13) and 3.7±1.0% (d21) oligodendrocytes (S1B Fig). Due to technical limitations (GFAP localization in astrocytic outgrowths and unavailability of a nuclear marker), astrocytes could not be automatically enumerated. The remaining population, namely total cells minus neurons and oligodendrocytes, comprising 21.5±4.1% (d13) and 22.2±4.4% (d21) of cells, was therefore considered to be composed of astrocytes (S1B Fig). The reliability of this enumeration procedure was confirmed by manual enumeration, as 22.8±5.6% (d13) and 32.7±5.0% (d21) of astrocytes were found using this method (S1C Fig). Thus, we confirmed that neurons and astrocytes were the major cell types in our co-cultures and showed that oligodendrocytes constituted less than 5% of the total cell population.

To characterize TBEV cellular tropism, we infected hNPC-derived neuronal/glial co-cultures and followed infection from 14 hours to 7 days. Cells were co-immunostained with antibodies directed against TBEV-E3 (infected cells) and βIII-tubulin or HuC/HuD (neurons), GFAP (astrocytes) and OLIG2 (oligodendrocytes). At 24 hpi, the 3 cell types were infected, as shown in fig. 2A. Viral envelope strongly accumulated in the perinuclear region of the cytoplasm in all cell types. The protein could also be evidenced in certain neurites and astrocyte outgrowths, albeit with a lower intensity (Fig. 2B). We then sought to determine whether the virus spread within each cellular subpopulation. We therefore quantified infection in each cell type, at different time points during the course of the study (Fig. 2C, 2D, 2E). The general profile of infection was similar in the 3 cell types, with an increase in the first days of infection up until a peak occurring at 48-72 hpi, followed by a decrease at 7 dpi, except for oligodendrocytes in which case no decrease was observed. Early in infection, at 14 hpi, a minority of cells were infected within each subset, namely 7.9±1.2% of neurons, 4.3±1.5% of astrocytes and 11.7±0.8% of oligodendrocytes. Later, however, at the peak of infection, the proportion of infected cells was high in neurons (55.2±3.8%) and oligodendrocytes (68.0±21.5%) but much lower in astrocytes (13.6±5.3%), revealing differential propagation of the virus within the three sub-population. Thus, whereas human neurons and oligodendrocytes were highly susceptible to TBEV infection, human astrocytes were more resistant.

**Figure 2.**
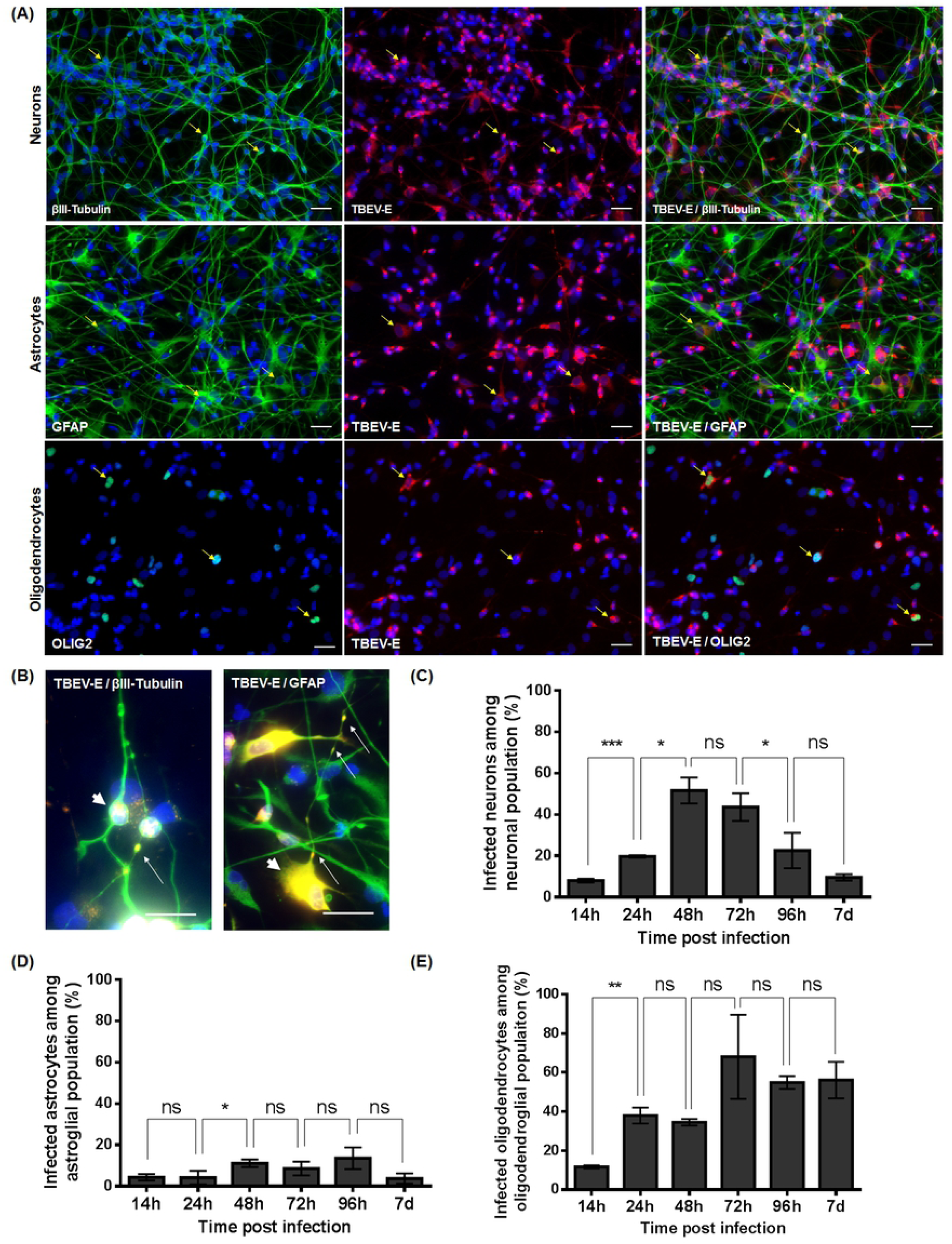
TBEV tropism for hNPCs-derived brain cells. HNPCs differentiated for 13 days were infected with TBEV-Hypr at MOI 10^-2^. (A) Immunofluorescence labeling of infected cells at 24 hpi. Antibodies against βIII-tubulin (neurones), GFAP (astrocytes) or OLIG2 (oligodendrocytes) (green), and TBEV-E3 (red) were used. Nuclei were stained with DAPI (blue). Yellow arrows show infected neurons, astrocytes and oligodendrocytes. Oligodendrocytes were recolored from grey to green. Scale bar=20 µm. (B) Higher magnification (digitally cropped) showing the viral envelope in perinuclear areas (arrowhead) and neurites and astrocytic outgrowths (arrows). Scale bar=20µm. (C-E) Percentage of infected cells based on immunofluorescence labeling during the course of infection for (C) neurons, (D) astrocytes and (E) oligodendrocytes. Results are representative of at least 2 independent experiments performed in triplicate. Data are expressed as the mean±SD. Statistical analysis was performed using a two-tailed unpaired t test with Graphpad Prism V6.0.1, ns=non-significant (p>0.05), *=p<0.05, **=p<0.01, ***=p<0.001.

### TBEV induces death of neurons and astrocytes

As hNPC-derived neuronal/glial cells were highly infected, we then sought to evaluate whether TBEV induced cellular damages. Cultures were infected and cells were fixed at several time points from 14 hpi to 14 dpi before immunostaining with antibodies specific for neuronal and glial cells, as previously described. We first examined the neuronal population. At 14 dpi, examination of HuC/HuD immunostaining revealed that TBEV-infected co-cultures were strongly depleted in neurons, as compared with their non-infected matched controls (Fig. 3A). Enumeration showed that neuronal survival was unaffected in the first days of infection (from 14 to 48 hpi), but confirmed that neuronal loss occurred as early as 72 hpi (25.1±5.4% loss) and steadily increased from this point on, reaching 72.0±10.3% at 14 dpi, the latest time point of our study (Fig. 3B). We then examined neuronal morphology based on βIII-tubulin immunostaining. At 7 dpi, a striking loss of neurites was observed in TBEV-infected cultures as compared with their non-infected matched controls (Fig. 3C). Quantification of total neurite length confirmed their loss not only at 7 dpi (76.1±21.6% decrease), but also at 72 hpi (62.0±22.6% decrease), whereas they were unaffected at an earlier time point (14 hpi) (Fig. 3D). Of note, whereas neuronal death became progressively more pronounced between 72 hpi and 7 dpi, neurite loss peaked as early as 72 hpi, suggesting that neurites alteration precedes neuronal death. Taken together, these results showed that TBEV infection strongly impaired neuronal survival in the co-cultures and, moreover, suggested that neurite alteration preceded neuronal death.

**Figure 3.**
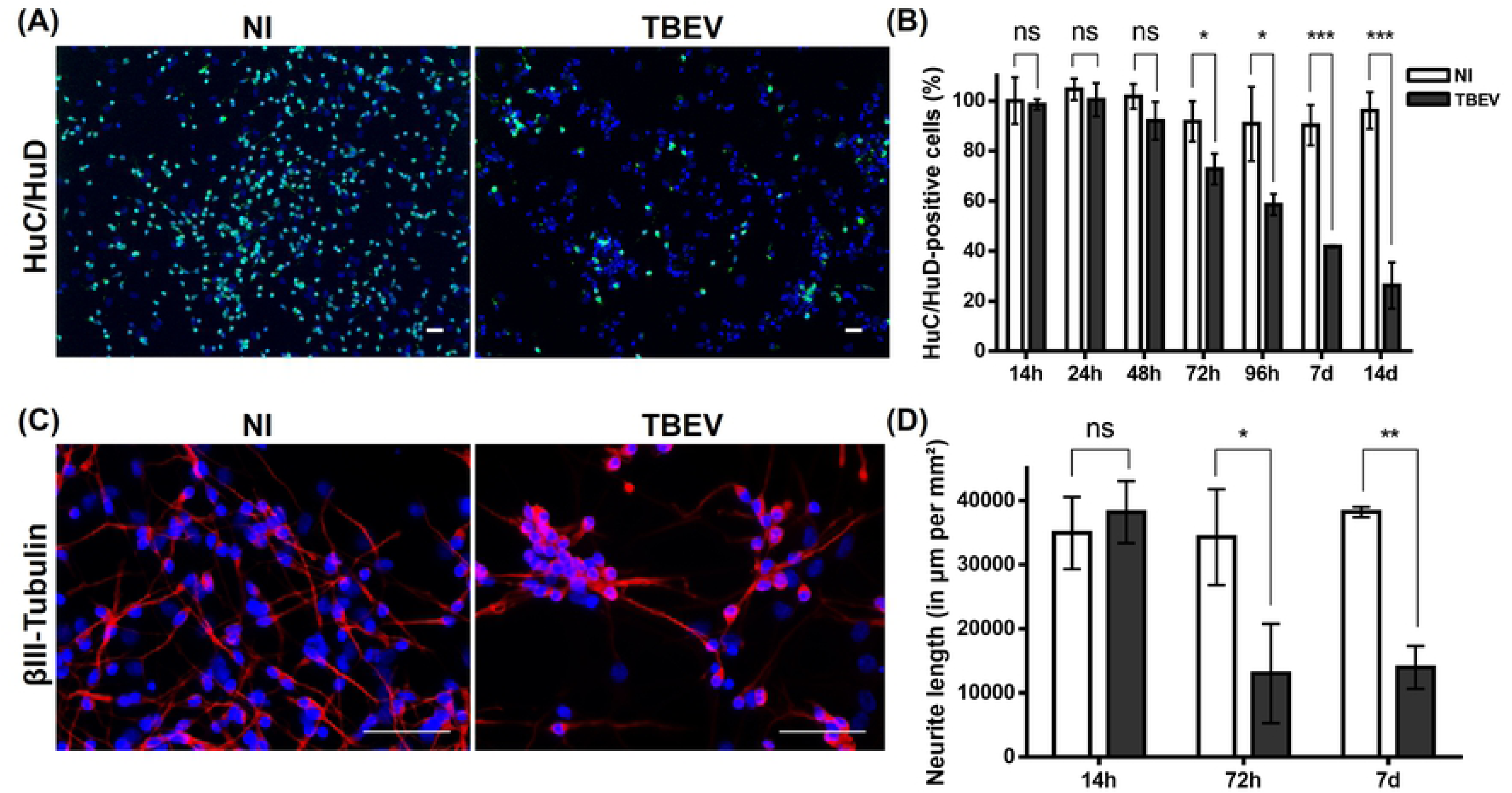
TBEV damages human neurons. HNPCs were differentiated for 13 days and infected with TBEV-Hypr at MOI 10^-2^. (A) Cells in non-infected (NI) and infected (TBEV) co-cultures were immunostained with an antibody directed against HuC/HuD (neurons, green) at 14 dpi. Nuclei were counterstained with DAPI. Scale bars=100µm. (B) Enumeration of HuC/HuD-positive cells using an ArrayScan Cellomics instrument. Normalization to non-infected HuC/HuD-positive cells at 14 hpi. (C) Cells in non-infected and infected cultures were immunostained with an antibody against βIII-tubulin (neurons, red) at 7 dpi. Note the paucity of neurites in TBEV-infected co-cultures. (D) Quantification of neurite network density (neurite length per mm²) using an ArrayScan Cellomics instrument. Results in (B) and (D) are expressed as the mean±SD and are representative of four and two independent experiments performed in triplicate, respectively. Statistical analysis was performed using a two-tailed unpaired t test with Graphpad Prism V6.0.1, ns=non-significant (p>0.05); *=p<0.05; **=p<0.01; ***=p<0.001.

We next evaluated whether glial cells were damaged. Examination of astrocytes immunostained with an antibody directed against GFAP at 7 dpi revealed hypertrophic cells in TBEV-infected cultures, as compared with their non-infected matched controls (Fig. 4A). This change in morphology is reminiscent of astrogliosis, a common feature of stressed astrocytes. Enumeration of GFAP-positive cells was then carried out at 24 hpi, 72 hpi and 7 dpi. Their number was not significantly altered at the earlier time points, 24 hpi and 72 hpi, but a decrease of 20.7±11.1% was observed at 7 dpi compared with non-infected matched controls (Fig. 4B). Thus, TBEV infection diminished survival of not only neurons but also astrocytes, although in a more moderate manner for the latter. By contrast, enumeration of OLIG-2-positive cells did not reveal a significant difference in oligodendrocyte number in TBEV-infected and non-infected cultures (Fig. 4C), showing that despite direct TBEV infection, survival of oligodendrocytes was unaffected. Taken together, our results demonstrated that subsets of hNPC-derived brain cells, that is, neurons, astrocytes and oligodendrocytes, were differentially affected by TBEV infection. In particular, neurons were highly susceptible as regards both infection and mortality, whereas astrocytes were more resistant. Oligodendrocytes were susceptible to infection, but their survival was unaffected.

**Figure 4.**
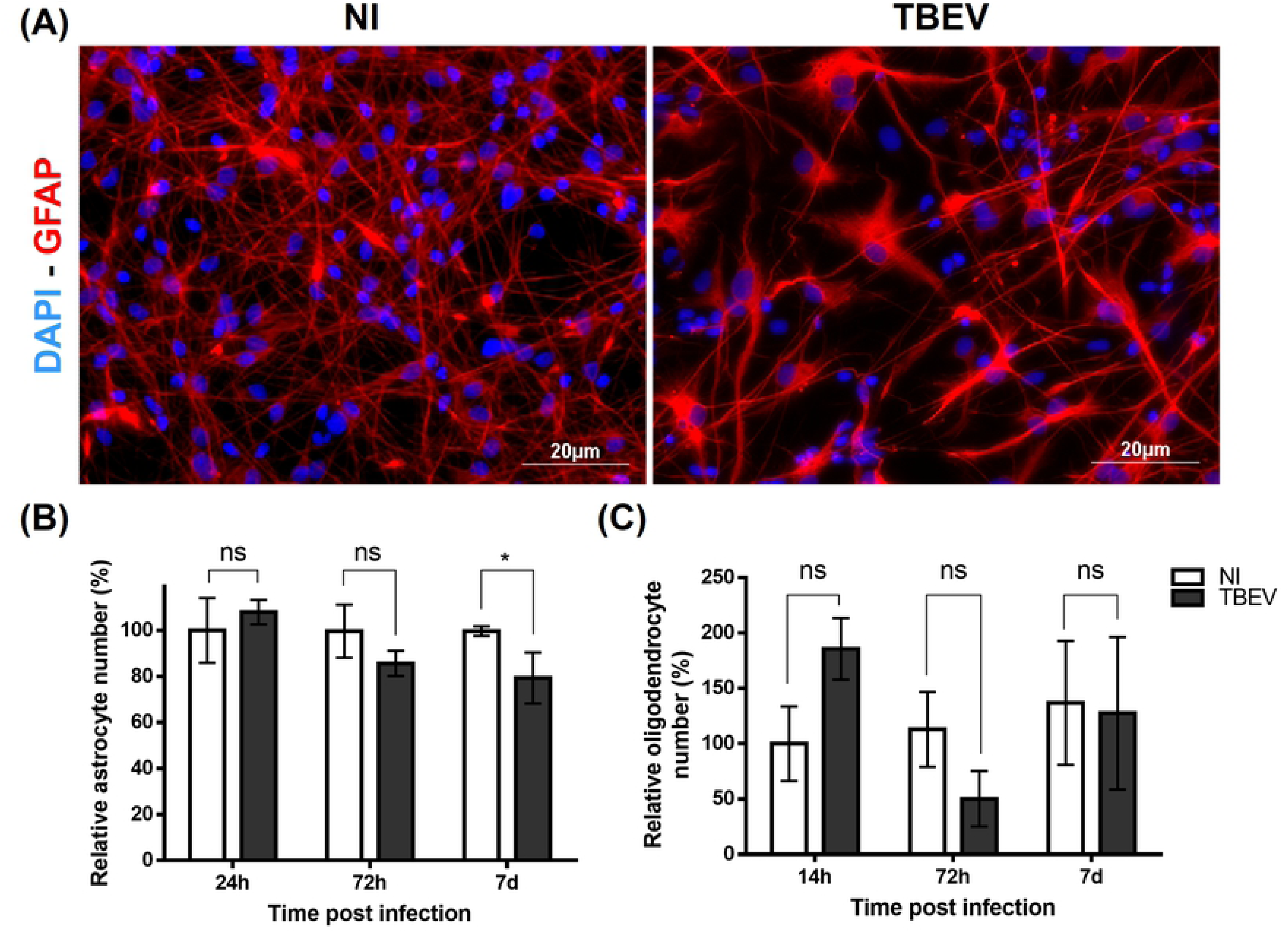
Impact of TBEV on human glial cells. HNPCs were differentiated for 13 days and infected with TBEV-Hypr at MOI 10^-2^. (A) Cells in non-infected (NI) and infected (TBEV) co-cultures were immunostained with an antibody directed against GFAP (astrocytes, red) at 7 dpi. Nuclei were counterstained with DAPI. Scale bars=20µm. (B) Manual enumeration of GFAP-positive cells using ImageJ software. Normalization was performed relative to non-infected GFAP-positive cells at 24 hpi. (C) Immunostained cells with OLIG2 antibody were enumerated automatically. Normalization was performed relative to non-infected OLIG2-positive cells at 24 hpi. The results are expressed as the mean±SD and are representative of two (oligodendrocytes) and three (astrocytes) independent experiments performed in triplicate. Statistical analysis was performed using a two-tailed unpaired t test with Graphpad Prism V6.0.1, ns=non-significant (p>0.05); *=p<0.05.

### Human NPC-derived neuronal/glial co-cultures develop a strong antiviral response to TBEV infection

When infected with virus, cells initiate an antiviral response that aims at controlling viral replication. In order to determine whether the human neuronal/glial cells used in our study had conserved the capacity to develop such a response upon TBEV infection, we analyzed the differential expression of 84 human genes involved in the antiviral response, using a PCR array approach. Transcripts from hNPC-derived neuronal/glial cells infected with TBEV for 24 h were pooled from biological triplicates and compared with their matched non-infected controls. The studied genes are shown in fig. 5A. After applying an arbitrary cut-off of 3-fold, 25 genes were shown to be significantly modulated in TBEV-infected cells, amongst which 22 genes were up-regulated and 3 were down-regulated (Fig. 5A). The former category included pathogen recognition receptors (PRRs), cytokines, including IFNβ, and ISGs. Overexpression of nine of these genes, 3 PRRs - IFIH1/MDA5 (Fig. 5B), DDX58/RIG-I (Fig. 5C) and TLR3 (Fig. 5D), 3 pro-inflammatory cytokines - CXCL10 (Fig. 5E), CCL5/RANTES (Fig. 5F) and CXCL11 (Fig. 5G) - and 3 ISGs - OAS2 (Fig. 5H), MX1 (Fig.5I) and ISG15 (Fig. 5J) - was confirmed using RT-qPCR. IFI6 (Fig. 5K), an additional ISG that was recently shown to protect cells from *Flavivirus* infection (30), was also shown to be overexpressed. For most of these genes, kinetic analyses further revealed that their expression was activated as early as 7 hpi and progressively increased during the course of infection up to 14 dpi, with the exception of pro-inflammatory cytokines whose expression abruptly decreased at 14 dpi (Fig. 5E-G). The latter, however, remained highly overexpressed, as compared with their matched non-infected controls. These data indicated that TBEV-infected hNPC-derived neuronal/glial cells had the capacity to respond to TBEV infection by developing a strong and lasting antiviral response.

**Figure 5.**
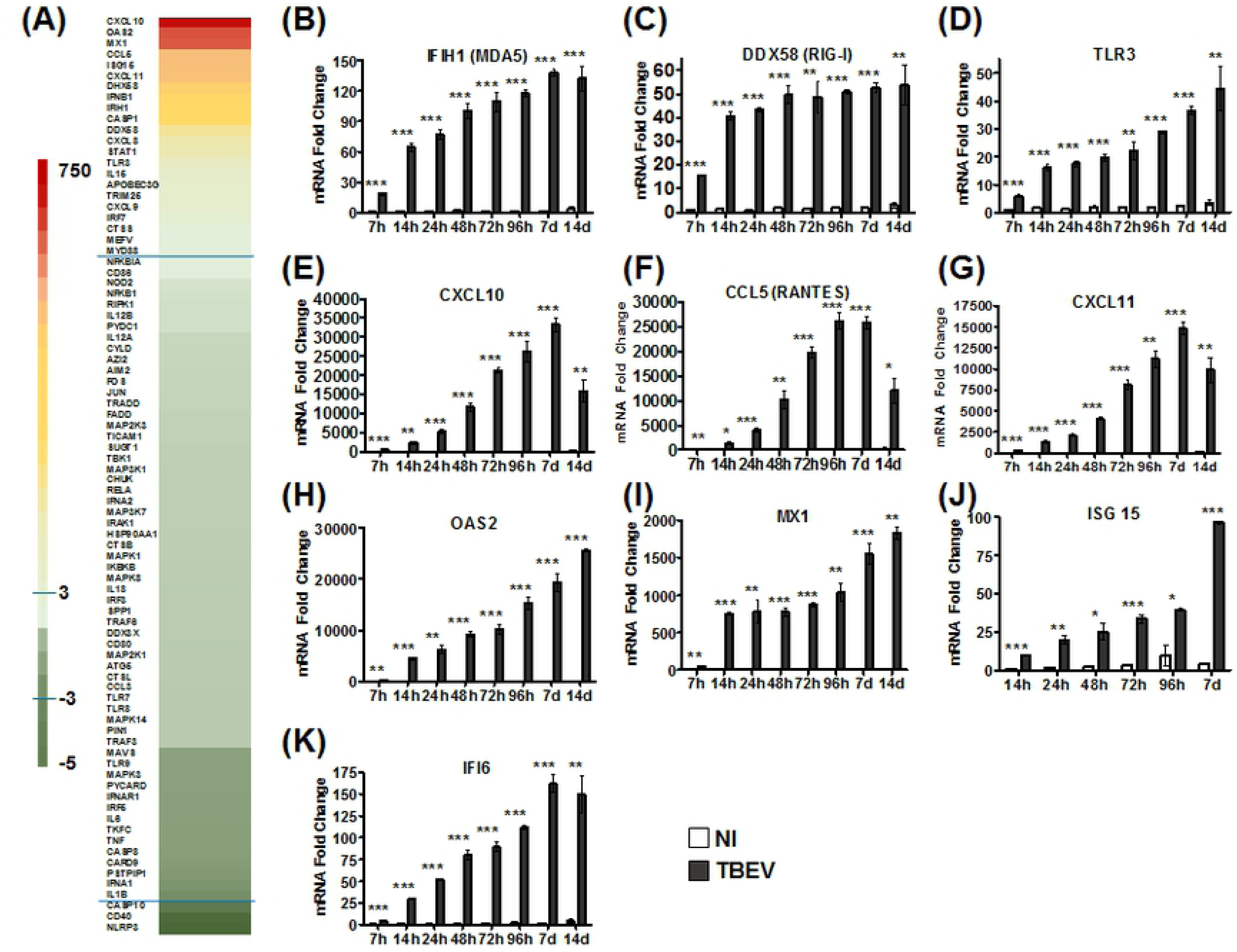
TBEV-induced antiviral response in hNPC-derived neuronal/glial cells. (A) TBEV-infected neuronal/glial cells and their matched NI controls were analyzed 24 hpi using an RT² Profiler PCR array specific for the human antiviral response. The heat map shows the differential expression of 84 analyzed human genes. The most highly up- and down-regulated genes are colored in red and dark green, respectively. The blue lines indicate the arbitrary cut-off of 3. Genes between the two lines are considered non-regulated. (B-K) RT-qPCR analyses of selected antiviral genes. Gene expression was normalized to *HPRT1* gene and the -2ΔΔCt method was used for relative quantification (normalization to non-infected cells at 7 hpi). Data are expressed as the mean±SD. Results are representative of one experiment performed on pooled triplicates (PCR array) or two independent experiments performed in triplicate (qPCRs). Statistical analysis was performed using a two-tailed unpaired t test with Graphpad Prism V6.0.1, ns=non-significant (p>0.05); *=p<0.05; **=p<0.01; ***=p<0.001.

### Differential antiviral response in human neurons and human astrocytes

Neurons and astrocytes are both known to participate in the antiviral response in the CNS (20, 31). As regards oligodendrocytes, little is known so far (32). Our results, showing high susceptibility of neurons but resistance of astrocytes to TBEV infection, led us to hypothesize that differences in their intrinsic capacity for antiviral defense might underlie their differential susceptibility. In order to test this hypothesis and decipher cell autonomous anti-TBEV innate immunity in the human CNS, we sought to obtain cultures enriched in neurons (henceforth called En-N) or astrocytes (henceforth called En-As) and to compare their antiviral response. Oligodendrocytes were not considered further in this study, as their low number in our cultures precluded enrichment. After differentiation of hNPCs for 13 days, neuronal/glial cells were trypsinized and either directly re-seeded (unsorted cultures henceforth called Uns-C) or enriched for neurons (En-N) or astrocytes (En-As). We showed that the splitting procedure did not alter the neuronal/glial co-cultures (Uns-C). Indeed, four days after re-seeding, phase-contrast microscopy of Uns-C revealed typical neuronal (small sized with neurites) and astroglial (larger, flat, with outgrowths) cells (Fig. 6A), as typically observed in non-trypsinized co-cultures (henceforth called Co-C cells). Cell type composition (74.1±4.1% neurons, 20.8±4.9% astrocytes and 5.1±1.2% oligodendrocytes) and basal expression of antiviral genes (analysis of 84 genes of the antiviral response) were unchanged in Uns-C as compared with Co-C cells (Fig. 6B and Fig. 6C, respectively). Enrichment of neurons and astrocytes was confirmed by phase-contrast microscopy (Fig. 6D and 6F, respectively) and cell enumeration, showing that the En-N population was composed of 94.1±0.4% neurons, 3.1±0.4% astrocytes and 2.8±0.2% oligodendrocytes (Fig. 6E) while the En-As population comprised 53.5±2.7% astrocytes, 35.7±2.8% neurons and 10.8±0.5% oligodendrocytes (Fig. 6 G).

**Figure 6.**
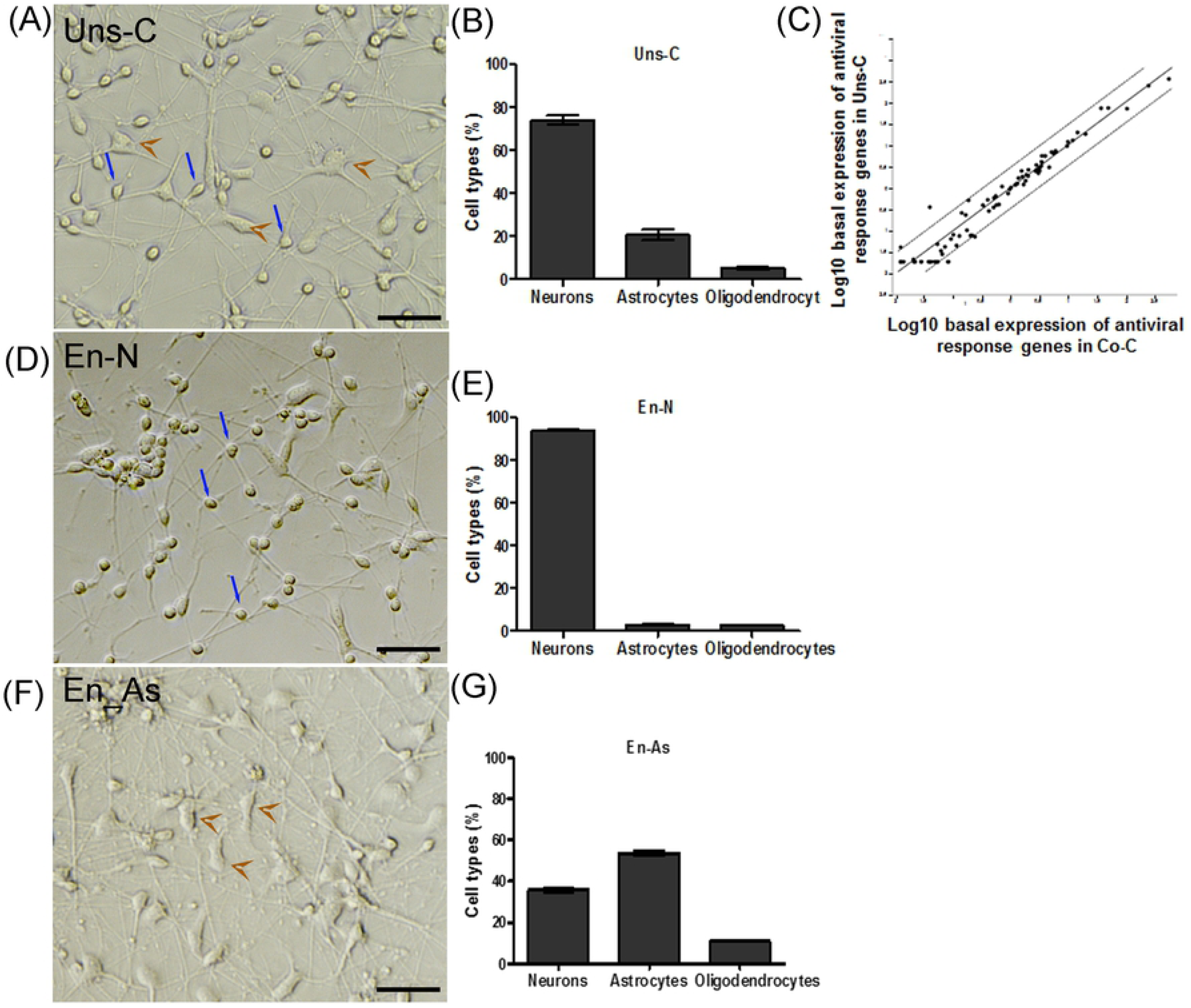
Enrichment of human neurons and astrocytes by magnetic-activated cell sorting. HNPC-derived neuronal/glial cells differentiated for 13 days were sorted using MACS technology. (A, D, F) Phase-contrast micrographs, showing Uns-C (A), En-N (D) and En-As (F), were acquired 96 h after re-seeding. Blue arrows indicate neurons and brown arrowheads indicate astrocytes. Scale bars = 50µm. (B, E, G). Enumeration of neurons, astrocytes and oligodendrocytes in Uns-C (B), En-N (E) and En-As (G) based on immunofluorescence staining (DAPI, antibodies against HuC/HuD and OLIG2) and using an ArrayScan Cellomics instrument. Data are representative of 4 independent experiments performed in triplicate. (C) Scatterplot of basal level of antiviral response genes in unsorted cells (Uns-C) compared with non-trypsinised cells (Co-C). Analysis was performed using an antiviral response PCR array. Genes along the black line have similar expression levels in the two cultures. Dotted lines represent an arbitrary cutoff of 3. Data are from a single experiment performed with pooled triplicates.

We then sought to determine whether distinct antiviral responses occurred in En-As, En-N and Uns-C upon TBEV infection, which would reflect differential antiviral responses in human neurons and astrocytes. Cells were infected for 24 h and the expression of 84 genes of the antiviral response was compared using the same PCR array as previously described. After application of the usual arbitrary cut-off of 3-fold, 20 genes in TBEV-infected Uns-C, 16 genes in TBEV-infected En-N and 21 genes in TBEV-infected En-As were shown to be significantly up-regulated, as compared with their non-infected matched controls (Fig. 7A). Among the set of over-expressed genes, which overlapped with that of non-trypsinized co-cultures, 13 were common to the three cultures (table 1) while others were specific for En-As or En-N cultures (8/21 and 3/16, respectively). Thus, these results showed that the antiviral program activated by TBEV was partially different in human neurons and astrocytes. Of note, for 12/13 of common genes, the magnitude of up-regulation was correlated to the percentage of astrocytes in the cultures. That is, it was much higher in En-As than in En-N and intermediary in Uns-C (Fig. 7A, table 1), showing that human astrocytes were capable of developing a stronger antiviral response to TBEV than human neurons. To validate the PCR array data and to gain further insight into the kinetics of expression of antiviral genes in each cell types, we performed RT-qPCR at 7, 24, and 72 hpi for 3 PRRs — IFIH1 (MDA5) (Fig. 7B), DDX58 (RIG-I) (Fig. 7C), and TLR3 (Fig. 7D) —, two ISGs — OAS2 (Fig. 7E) and MX1 (Fig. 7F) —, and one pro-inflammatory cytokine, CXCL10 (Fig. 7G). Because of their well-known anti-flavivirus activity, the ISGs IFI6 (Fig. 7H) and RSAD2 (viperin) (Fig. 7I) were also studied. In confirmation of the PCR array data, all of these genes were significantly more overexpressed in En-As than in En-N, at both 24 and 72 hpi. In addition, at 72 hpi, gene expression in En-N was either maintained (RIG-I, TLR3, OAS2, viperin, CXCL10) or decreased (MDA5, MX1, IFI6) while it was either maintained (MDA5, TLR3, MX1, viperin) or increased (RIG-I, OAS2, CXCL10, IFI6) in En-As, showing that the duration of antiviral responses was shorter in human neurons than astrocytes. The induction of an astrocyte-specific antiviral program, as suggested by the selective overexpression of 8 genes in En-As (Fig. 7A, Table 1), was further confirmed by the results of RT-qPCR. Not only TLR3 but also viperin were amongst those genes, as upregulation was observed in En-As at 24 hpi and 72 hpi but not in En-N at either time point (Fig. 7D and Fig. 7I). Thus, taken together, these results show that TBEV infection induces an antiviral response in human neurons and astrocytes that is characterized by activation of an overlapping set of genes. This anti-viral program, however, was of greater intensity and longer duration in human astrocytes than in human neurons. The anti-viral programs of the two cell types were also notably distinct, as exemplified by selective overexpression of the gene encoding viperin, well-known for its anti-TBEV activity (33, 34), in human astrocytes. In sum, our results revealed a stronger and broader antiviral response in human astrocytes than in human neurons, in keeping with their differential susceptibility to TBEV infection, astrocytes being more resistant and neurons more susceptible.

**Figure 7.**
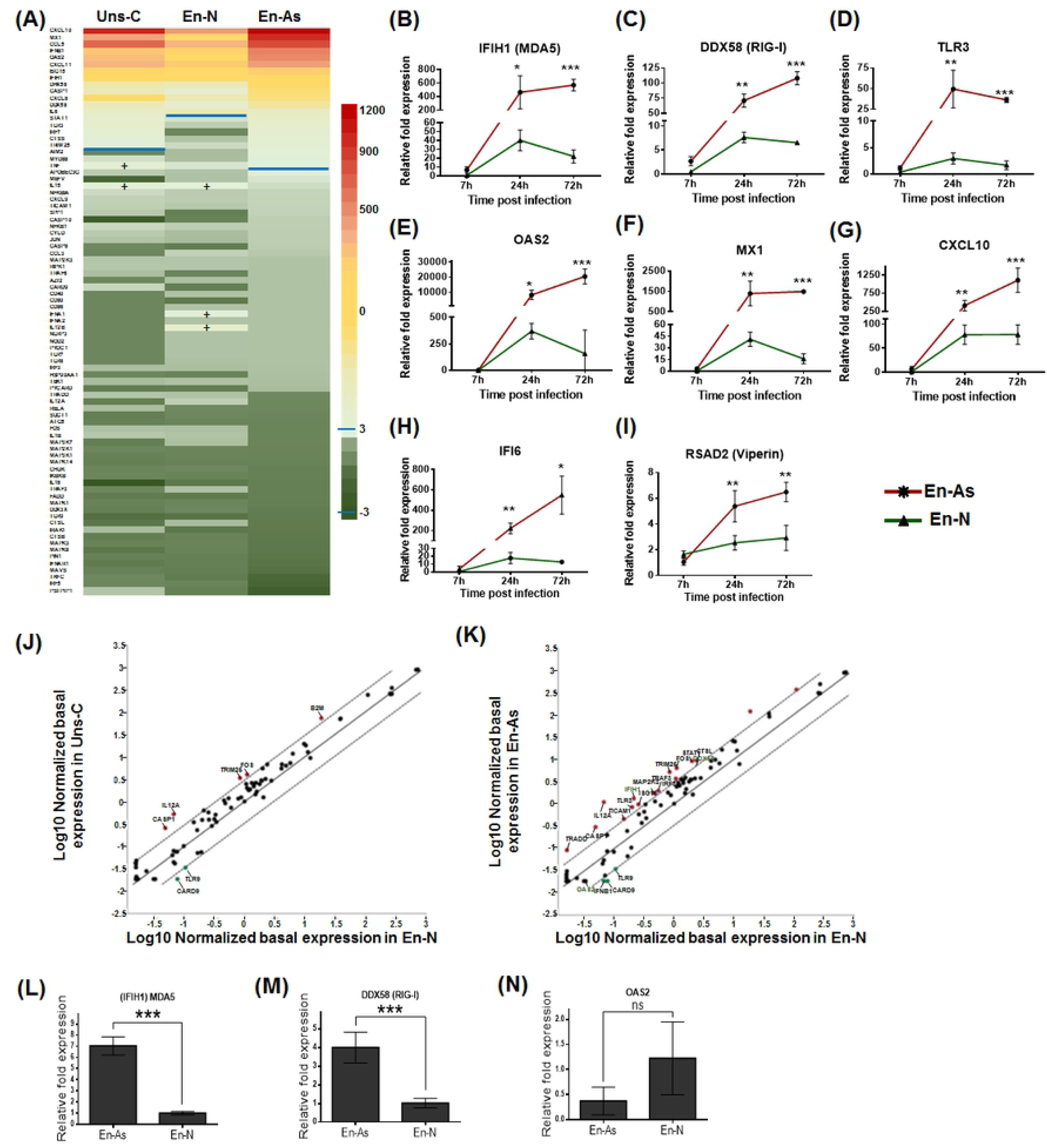
Antiviral response in enriched neurons, enriched astrocytes and unsorted cells. (A-I) Expression of antiviral genes upon TBEV infection. (A) TBEV-infected Uns-C, En-N and En-As and their matched non-infected controls were analyzed 24 hpi using an RT^2^ profiler PCR array specific for the human antiviral response. Heat map showing the 84 human genes analyzed and their differential expression. Color code and blue line are as in figure 5. Note that genes noted (+) are above the cut-off of 3. (B-I) RT-qPCR analyses of selected antiviral response genes in En-As (red) and En-N (green). (J-N) Basal expression of antiviral genes. (J, K) Scatterplots of basal expression levels of antiviral response genes in enriched neurons (En-N) compared with that of unsorted cells (Uns-C) (J) or enriched astrocytes (En-As) (K). Analysis was performed using an antiviral response PCR array. (L-N) RT-qPCR analysis of the basal expression of IFIH1/MDA5 (L), DDX58/RIG-I (M), and OAS2 (N) genes in En-N and En-As. Gene expression was normalized to HPRT1 and the -2ΔΔCt method was used for relative quantification (normalization to non-infected En-N, at 7hpi for B-I). The results are expressed as the mean±SD. Data are representative of two independent experiments performed in triplicate (B-I, L-N) and one experiment performed with pooled triplicates (J, K). Statistical analyses comparing En-As and En-N were performed with Graphpad Prism V6.0.1 using a two-tailed unpaired t test, ns=non-significant (p>0.05); *=p<0.05; **=p<0.01; ***=p<0.001.

**Table 1.**
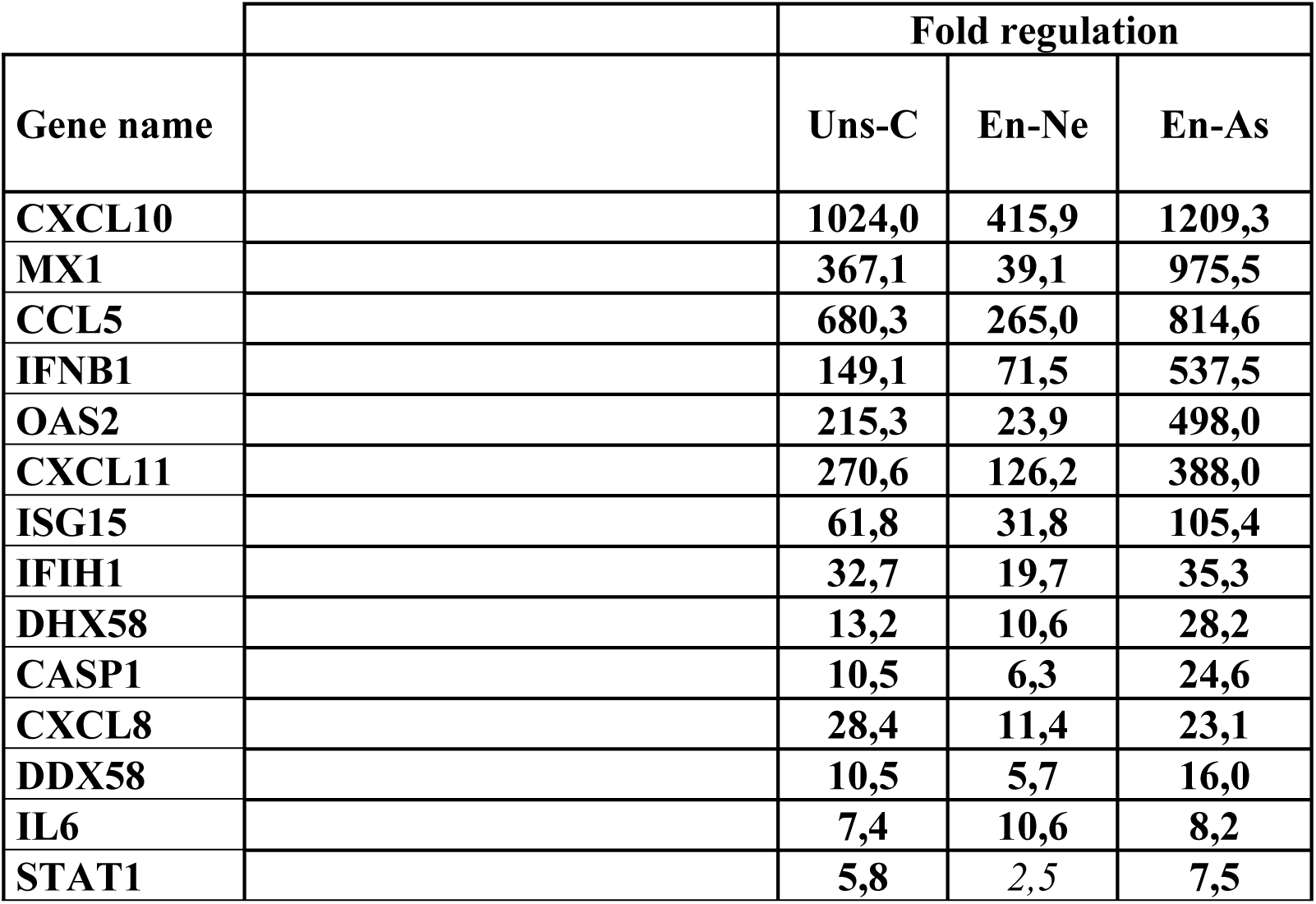
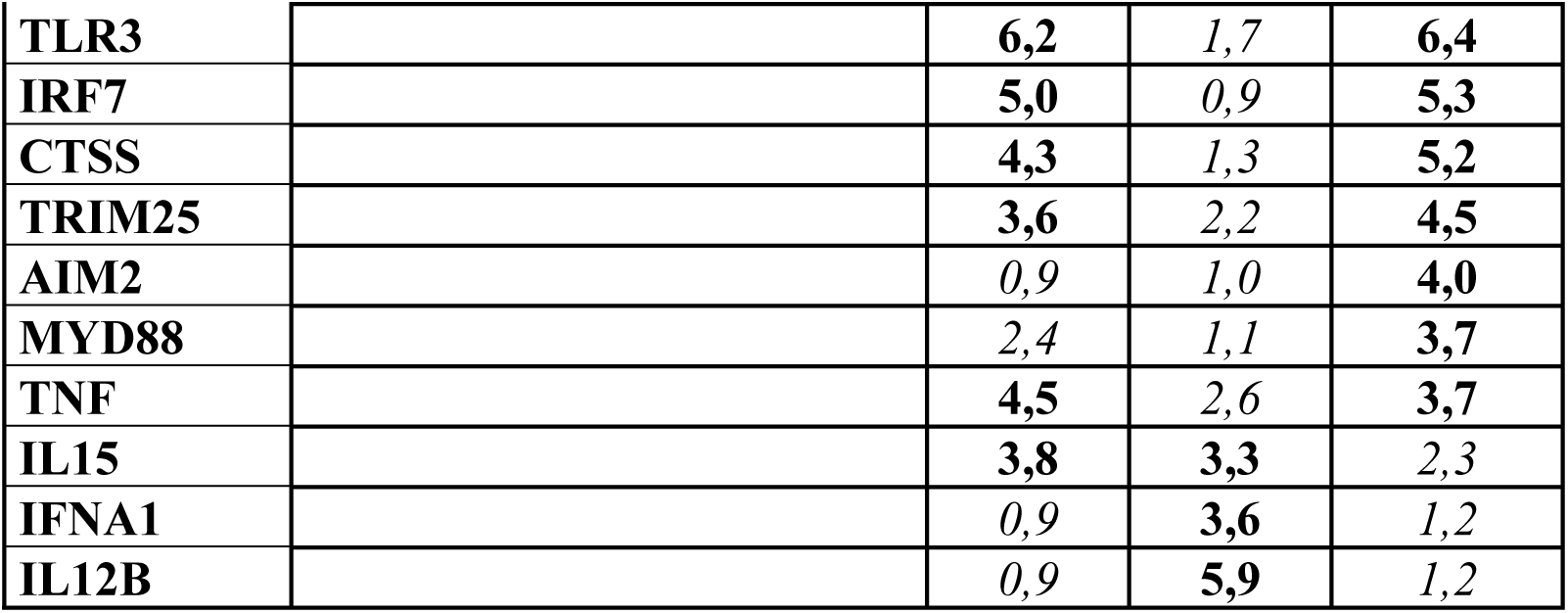
Differential expression of genes involved in the human antiviral response in unsorted cultures (Uns-C), enriched neurons (En-N) and enriched astrocytes (En-As). Up and down-regulated genes appear in bold, after application of a cut-off of 3.

| Gene name |  | Fold regulation |  |  |
| --- | --- | --- | --- | --- |
|  |  | Uns-C | En-Ne | En-As |
| **CXCL10** |  | **1024,0** | **415,9** | **1209,3** |
| **MX1** |  | **367,1** | **39,1** | **975,5** |
| **CCL5** |  | **680,3** | **265,0** | **814,6** |
| **IFNB1** |  | **149,1** | **71,5** | **537,5** |
| **OAS2** |  | **215,3** | **23,9** | **498,0** |
| **CXCL11** |  | **270,6** | **126,2** | **388,0** |
| **ISG15** |  | **61,8** | **31,8** | **105,4** |
| **IFIH1** |  | **32,7** | **19,7** | **35,3** |
| **DHX58** |  | **13,2** | **10,6** | **28,2** |
| **CASP1** |  | **10,5** | **6,3** | **24,6** |
| **CXCL8** |  | **28,4** | **11,4** | **23,1** |
| **DDX58** |  | **10,5** | **5,7** | **16,0** |
| **IL6** |  | **7,4** | **10,6** | **8,2** |
| **STAT1** |  | **5,8** | **2,5** | **7,5** |
| **TLR3** |  | **6,2** | *1,7* | **6,4** |
| **IRF7** |  | **5,0** | *0,9* | **5,3** |
| **CTSS** |  | **4,3** | *1,3* | **5,2** |
| **TRIM25** |  | **3,6** | *2,2* | **4,5** |
| **AIM2** |  | *0,9* | *1,0* | **4,0** |
| **MYD88** |  | *2,4* | *1,1* | **3,7** |
| **TNF** |  | **4,5** | *2,6* | **3,7** |
| **IL15** |  | **3,8** | **3,3** | *2,3* |
| **IFNA1** |  | *0,9* | **3,6** | *1,2* |
| **IL12B** |  | *0,9* | **5,9** | *1,2* |

Differential expression of antiviral genes in human neurons and astrocytes following TBEV infection may reflect differential baseline expression in the two cell types. To test this hypothesis, non-infected En-N, Uns-C and En-As cells were cultured for 4 days and transcripts from 3 biological samples in each condition were pooled and compared using the human antiviral response PCR array. Antiviral gene expression in En-N was compared with that of Uns-C (Fig. 7J) and En-As (Fig. 7K). Although in both cases, most of the immunity-related genes were not differentially expressed to a significant extent (above the 3-fold threshold recommended by the manufacturer), we observed that the general tendency was to a slight overexpression in astrocytes, since the number of significantly overexpressed genes increased as the percentage of astrocytes increased in the culture, from Uns-C to En-A. Five/84 genes were, indeed, overexpressed in Uns-C compared with En-N (Fig. 6J), while their number rose to 14/84 genes when En-As were compared with En-N (Fig. 6K). By contrast, very few of these genes (3/84) were overexpressed in human neurons, and then only to a modest extent. To validate these results, differential expression of 3 selected genes, 2 PRRs (MDA5 and RIG-I) and 1 ISG (OAS2), was further addressed by RT-qPCR. Significant overexpression of MDA5 (Fig. 7L) and RIG-I (Fig. 7M) in En-As as compared with En-N was confirmed. By contrast, but still in agreement with the PCR array data, no significant difference was observed for the OAS2 gene (Fig. 7N). These results thus showed that the basal level of expression of certain antiviral genes was higher, albeit slightly, in human astrocytes than in human neurons.

### Human astrocytes protected human neurons from TBEV infection and TBEV-induced damage

Next, we wondered whether human astrocytes might participate in neuronal defense. We reasoned that, if this were the case, neurons would be more sensitive to TBEV infection in cultures depleted of astrocytes. We therefore compared neuronal susceptibility and vulnerability to TBEV infection in Uns-C and En-N, composed of 20.8±4.9% and 3.1±0.4% of astrocytes, respectively. Uns-C and En-N were infected for 24h and the percentage of TBEV-infected neurons within the total neuronal population was quantified based on βIII-tubulin and TBEV-E3 immunostaining. A 30% increase in infected cells was observed in En-N compared with Uns-C (Fig. 8A), showing that, in the absence of astrocytes, neurons were more sensitive to TBEV infection. At that time, neuronal survival was altered neither in TBEV-infected Uns-C nor in TBEV-infected En-N, as revealed by observation of βIII-tubulin immunostaining (Fig. 8B) and enumeration of neurons (S2 Fig.). At 72 hpi, however, a more dramatic alteration of neuronal morphology was observed in TBEV-infected En-N, as compact clusters, characteristic of intense neuronal death, were formed (Fig. 8B). Due to these clusters, it was not possible to enumerate the neurons, but our results clearly showed that, at 72 hpi, neurons were more dramatically affected in cultures deprived of astrocytes. Thus, taken together, these results showed that, upon TBEV infection, the presence of astrocytes was protective for human neurons.

**Figure 8.**
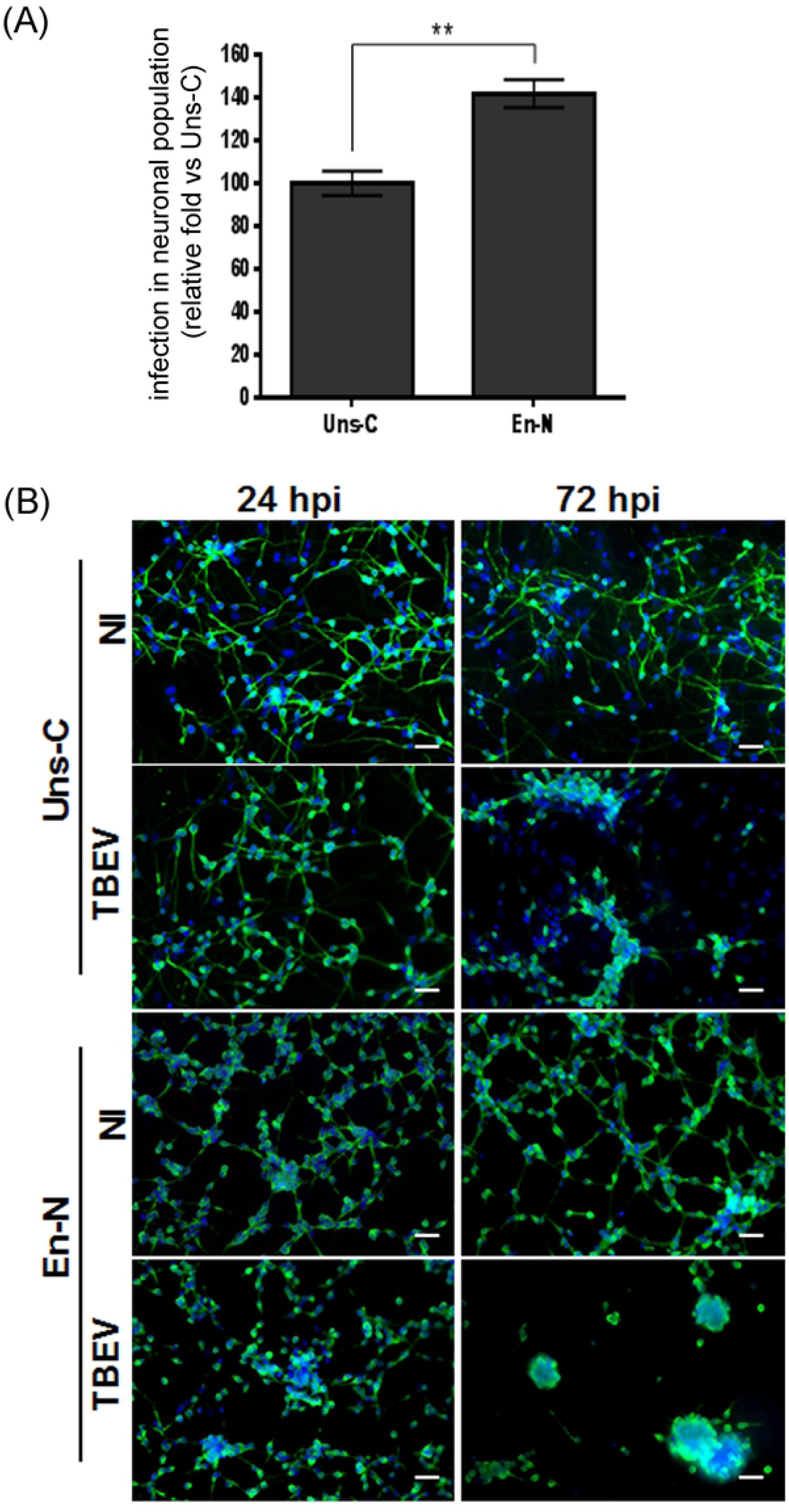
Human astrocytes protect neurons from TBEV infection. Unsorted cells (Uns-C) and enriched neurons (En-N) were infected with TBEV and co-immunostained using anti-βIII-tubulin and anti-TBEV-E3 antibodies. (A) Infected neurons among the neuronal population. Manual enumeration. The data is displayed as relative infection to Uns-C at 24 hpi. Data are expressed as the mean±SD. (B) Immunofluorescence staining of neurons (green). Nuclei were stained with DAPI (blue). Scale bar=20µm. Results are representative of two independent experiments performed in triplicate. Statistical analyses were performed with Graphpad Prism V6.0.1 using a two-tailed unpaired t test, **=p<0.01.

## Discussion

Despite its importance in human health, TBEV-induced neuropathogenesis is still poorly understood. So far, most studies have used either *in vitro* or *in vivo* rodent models. Whereas these models have advanced understanding, extrapolation to human neuropathogenesis may not always be relevant, as cellular responses display profound differences between species (25,26, 35). Here we used neuronal/glial cultures derived from human fetal neural progenitor cells as a more accurate *in vitro* model to study anti-TBEV innate immunity and its relation to tropism and neuropathogenesis in the human brain. We developed a new *in vitro* model that mimics major hallmarks of TBEV infection in the human brain, namely, neuronal tropism, neuronal death and astrogliosis, thereby providing a unique and highly relevant pathological model for studying TBEV-induced neuropathogenesis. Moreover, we revealed that a cell-type-specific innate antiviral state in human neurons and astrocytes correlates with their differential susceptibility and vulnerability to TBEV, which strongly suggests that the innate antiviral response shapes TBEV tropism for human brain cells.

Understanding viral tropism is critical for understanding virus-induced neuropathogenesis. Some viruses, like Zika virus, preferentially infect neural progenitors (36, 37) whereas human immunodeficiency virus has a strong affinity for microglial cells (38), JC virus for astrocytes (39) and the JHM strain of mouse hepatitis virus (JHMV) for oligodendrocytes (40). Flaviviruses such as TBEV, WNV and JEV have a preferential tropism for neurons, a feature that has been observed in human patients (12,41,42), as well as in rodent models (43, 44). Using human neuronal/glial cultures, we reproduced the preferential neuronal tropism that is observed *in vivo* in showing a high percentage of infected neurons (approximately 55% at the peak of infection) together with a low percentage of infected astrocytes (less than 15 %). The limited capacity of the virus to infect astrocytes, as observed in our cultures, as well as in monocultures of rat or human astrocytes (45–47), may explain the lack of detection of infected astrocytes in post-mortem brain tissues from patients with tick-borne encephalitis (12), or their rare detection in Langat virus-infected mice (44), as infrequently infected cells are likely to escape detection. Similar observations have been made for other neurotropic viruses (48–50). Unexpectedly, we also observed infection of oligodendrocytes in the human neuronal/glial cultures, a finding that has never been reported previously. The *in vivo* significance of this observation is at present unknown, especially because the degree of maturation of oligodendrocytes in our culture is undefined, the OLIG2 antibody recognizing mature and immature cells indiscriminately (51). However, this should be kept in mind for future examination of brain tissues from infected patients. As these cells represent about 5 % of the total cell population in our cultures, we believe them to have a minor impact *in vitro* and, as we could not enrich them, they were not further considered and we confined our study to neurons and astrocytes. We questioned the reasons that may explain difference in TBEV tropism for these two cell types. This may be due to differential expression of cellular factors that are necessary for establishing a full viral cycle (entry and post-entry events), but the similar percentage of infected neurons and astrocytes that we observed in the first 14 hours following TBEV infection does not lend support for this hypothesis. An alternative hypothesis would be differential capacity of the cell types to develop a protective antiviral response. The innate immune response, a major component of the antiviral response, has indeed been proven to be critically important in restricting infection by neurotropic viruses (52–54) and in determining TBEV tropism in different brain areas in murine models (44, 55). Also, studies using rodent models have shown that distinct brain cell types develop different antiviral states. Microglia and astrocytes, for example, which were initially considered to be the sole sentinels that respond to microbial infection within the brain (56, 57) have been shown to behave differently, as microglia developed a more robust response than astrocytes to TLR7 activation (58). Neurons, long considered to be merely passive targets, are now known to participate in the antiviral response and viral restriction (19–21,59) and in humans, neurons and astrocytes have been shown to produce and respond to IFNα/β. Despite a general assumption that astrocytes are more important players in antiviral response than neurons, the relative contribution of each cell types has, however, not been formally demonstrated, as direct comparison has never been made, whether in animal or in human *in vitro* models. Here we provided the first evidence that human neurons derived from fetal neural progenitor cells possess all of the necessary machinery to mount a cell-intrinsic antiviral response against TBEV, as they up-regulated IFNβ, ISGs and pro-inflammatory cytokine mRNAs upon TBEV infection. We also demonstrated for the first time that human neurons and astrocytes differ in their capacity to mount an anti-TBEV response. Differences were indeed observed in the repertoire of the antiviral program that is activated in the two cell types upon TBEV infection, with certain genes over-expressed in astrocytes but not in neurons, such as the RSAD2 gene (encoding viperin), an ISG that has been shown to be highly important for controlling TBEV (33, 34) and other flaviviruses (55) in the rodent’s CNS. Quantitative differences were also observed as transcripts encoding PRRs and genes of IFN signaling were over-expressed with different magnitudes in human neurons and astrocytes, with a stronger and more durable response in astrocytes than in neurons. Our results thus show that the cell-type specific anti-TBEV response is correlated with the susceptibility of human neurons and astrocytes to TBEV, which strongly suggests that innate antiviral response is responsible for shaping TBEV tropism in human brain cells. Whether the high neuronal susceptibility to TBEV infection is due to a weaker general antiviral program in human neurons, involving multiple components of the IFN response, from the PRRs to ISGs, or rather to lower expression of specific ISGs, such as the RSAD2 or IFI6 genes, that are dedicated to the control of flaviviruses (30), remains to be elucidated. It also remains to be determined whether the antiviral response of neurons is weak in response to infection by any neurotropic viruses, or only by certain viruses, such as flaviviruses, or whether it is specific to TBEV infection. Our observation that the baseline expression of certain antiviral genes is lower in neurons than in astrocytes may argue in favor of the first possibility, a hypothesis that should be addressed in future studies. In contrast to our results, TBEV infection in the human neuronal DAOY cells, a human neuroblastoma, led to overexpression of the RSAD2 gene (60), a discrepancy that may be due to the use of non-physiological, immortalized cells in the study from Selinger et al. (2017). Of note, our results showed that neuronal tropism of TBEV does not depend only on the cell-specific antiviral response in human neurons, as the presence of astrocytes in the culture limited their infection and favored their survival. As it was previously reported that murine astrocytes infected with TBEV were protective to neurons through IFN signaling (45), and as we showed that IFNβ was highly up-regulated by human astrocytes upon TBEV infection, we speculate that IFNβ produced by astrocytes acts in a paracrine manner to restrict neuronal infection in our human neuronal/glial co-cultures.

Viral tropism and pathogenesis are intimately linked, but how the former governs the latter in the human CNS, during TBEV infection, is incompletely understood. TBEV preferentially infects and kills the neurons (12), a highly dramatic event, as neurons have a very poor capacity to regenerate. Neuronal death may occur either in direct or indirect manners, such as, in the latter case, by inducing secretion of neurotoxic proteins by resident glial cells or recruitment of peripheral inflammatory cells to the brain parenchyma (11, 14). Involvement of T cells in neuronal death cannot be explored in our model. However, it has to be noted that chemokines such as CXCL10, CCL5 and CXCL11, which have been shown to be over-expressed in the cerebrospinal fluid of human patients infected with TBEV (61–63), are highly up-regulated by both human neurons and astrocytes, revealing that the two cell types may participate in chemo-attraction of T cells into TBEV-infected human brain parenchyma. Similar to the observation made for PRRs and ISGs, their overexpression was, however, stronger in astrocytes than in neurons, showing that astrocytes may be a major player in this process as well. As for neuronal death due to direct infection by TBEV, it has not yet been demonstrated, although ultrastructural changes in response to TBEV infection have been observed (64, 65). In our human neuronal/glial and enriched neuron cultures, neuronal death occurred in the absence of peripheral cells and was associated with infection of a large proportion of neurons, showing that the virus is directly responsible for their death. This is likely to play an important role in the human brain upon infection with TBEV and most probably other flaviviruses, since similar conclusions have been drawn for West Nile virus in studies performed in primary murine neurons (66). Reactive astrocytes, however, may also influence neuronal death. Astrogliosis, indeed, occurs following brain trauma of diverse etiology (67), including infection by TBEV or the related Langat virus infection (12), and may be either beneficial or detrimental to neurons (68). In human neuronal/glial cultures, we have observed that astrocytes were hypertrophic, a classical feature of reactive astrocytes and, when neurons were deprived of astrocytes, they were more sensitive to TBEV infection, showing that astrocytes exerted, under these conditions, a neuroprotective effect, possibly via restriction of neuronal infection as previously discussed. Thus, the intensity of neuronal death depends not only on direct infection of neurons but also on the indirect effect mediated by reactive astrocytes. Proliferation sometimes accompanied astrocyte hypertrophy in astrogliosis. Surprisingly, in our experiments, we observed a decrease in the number of astrocytes by 7 days following infection, which strongly suggested that TBEV induced astrocytic death. This is in apparent contradiction with previous work showing that astrocyte viability was unaffected in rodent and human astrocyte monocultures (46, 47). The difference in our results may be due to differential conditions of infection, such as viral strain or infectious dose or to the presence of neurons in our co-cultures, which may lead to astrocyte death by an unknown mechanism. Of note, infection with TBEV is not always correlated to cell death, since as many as 60 % of oligodendrocytes were infected without impact on their viability. Why certain cell types die upon TBEV infection while others do not remains to be understood, as do the molecular pathways that lead to cell death. Interestingly, our findings suggest that neuronal death may result from axonal pathology and retrograde degeneration. Indeed, we showed that loss of neurites preceded the disappearance of neuronal cell bodies, an observation that is in agreement with previous work in which accumulation of viral protein in neuritic extensions and dendritic degeneration due to local replication of TBEV was evidenced in murine neurons (64). The relative contribution of axonal degeneration, which is known to play a pathogenic role in rabies virus infection (69) and in some neurodegenerative diseases (70), to TBEV-induced neuronal death need to be defined in future studies.

Until recently, the lack of relevant *in vitro* models virtually precluded meaningful study of viral pathogenesis in the human brain. This obstacle has been overcome by the development of methodologies providing an unlimited source of human neural cell types that can be used for disease modeling. In this study, we set up a new, complex and highly relevant *in vitro* model that mimics the major events of TBEV infection in the human brain. Using this model, we evidenced differential innate immune responses in human neurons and astrocytes that contribute to shaping TBEV tropism and neuro-pathogenesis. Based on our results, we propose a model for interactions between TBEV and human brain cells that is represented in fig 9. Our study thus advances understanding of the mechanisms involved in TBEV-induced damage of the human brain and provides a pathological model that can be used in the future to provide greater knowledge as well as to develop new therapies by screening for antiviral or neuroprotective drugs.

**Figure 9.**
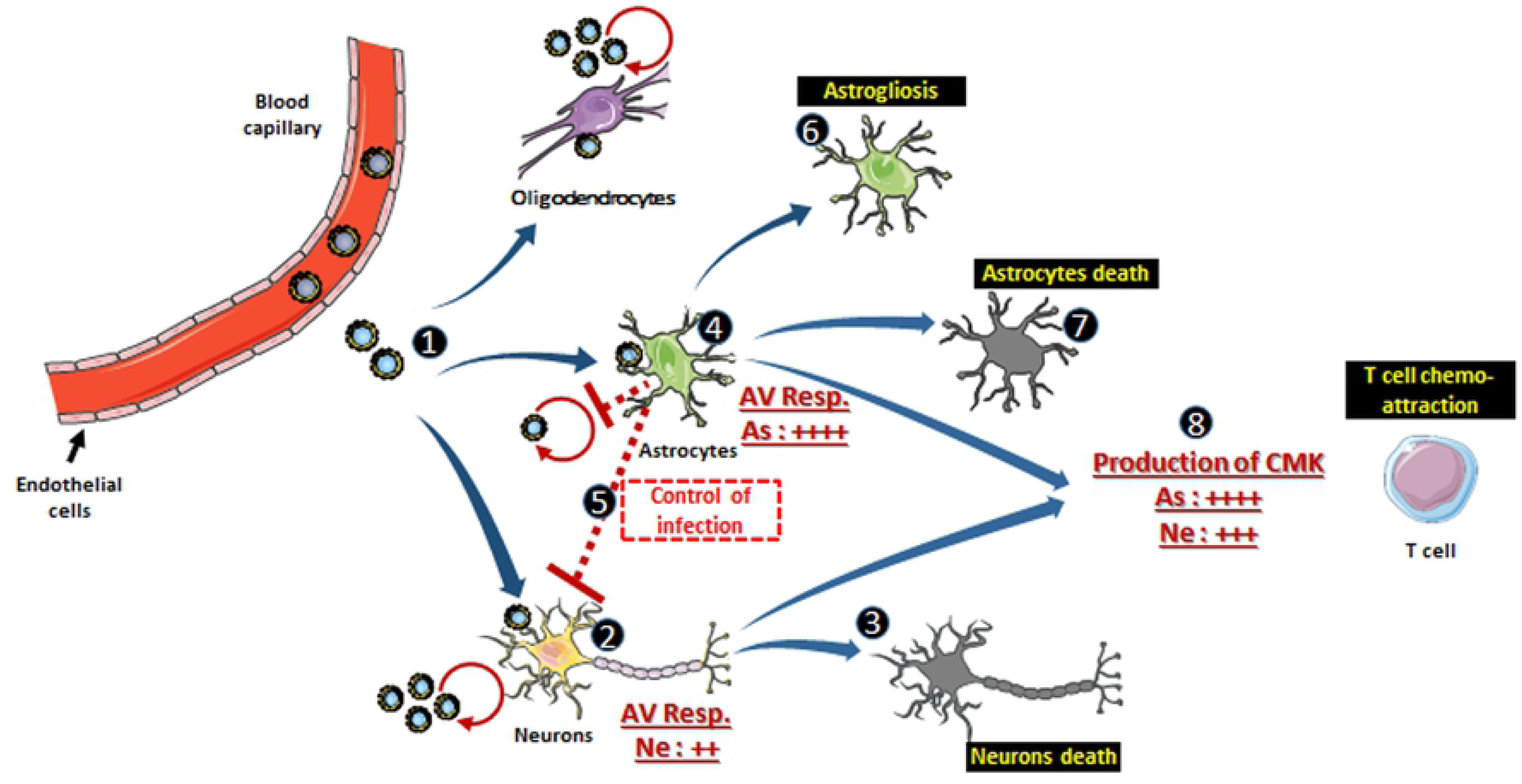
Proposed model of interactions between TBEV and human brain cells. In the human brain parenchyma, TBEV infects neurons, astrocytes and possibly oligodendrocytes (1). Both neurons and astrocytes develop an antiviral response. In neurons, it is insufficient to afford protection (2) and poorly controlled infection induces neuronal death in a direct manner (3). Astrocytes are infected but control infection, owing to their strong antiviral response (4), which may also be beneficial to neurons (5). Astrocytes enter a reactive stage (6) and some of them die (7). Both neurons and astrocytes overexpressed a high level of chemokines involved in chemo-attraction of T cells in the brain parenchyma (8), although astrocytes are stronger producers. Figure was created using Servier Medical Art available on www.servier.com. As, astrocytes. AV Resp, antiviral response. CMK, Chemiokines. Ne, neurons.

## Material and methods

### Ethics statement

Human fetuses were obtained after legal abortion with written informed consent from the patient. The procedure for the procurement and use of human fetal central nervous system tissue was approved and monitored by the “Comité Consultatif de Protection des Personnes dans la Recherche Biomédicale” of Henri Mondor Hospital, France. The cells are declared at the “Centre de Ressources Biologiques“ of the University Hospital in Angers BB-0033-00038 with reference numbers at the Research Ministry: declaration N° DC-2011-1467; authorization N° AC-2012-1507.

### Culture of human neural progenitor cells

Human neural progenitor cells (hNPCs) were prepared and cultured as previously described in (28, 29)

### Neuronal and glial differentiation

hNPCs were seeded on matrigel-coated plates at a density of 30 000 cells/cm². Differentiation to a mixed population of neuronal and glial cells was induced 24 h after plating by replacing N2A medium with 1:1 N2A and NBC media (N2A: advanced Dulbecco’smodified Eagel medium-F12 supplemented with 2mM L-glutamine, 0.1mg/ml apotransferrin, 25µg/ml insulin and 6.3ng/ml progesterone. NBC: Neurobasal medium supplemented with 2mM L-glutamine and B27 without vitamin A 1X - Invitrogen, Life Technologies) and withdrawing EGF (TEBU, France) and bFGF (TEBU, France). Differentiation conditions were maintained for 13 days with medium replacement twice a week, prior to infection. Twenty-four-well plates (IBIDI, #82406) were used for fluorescent immunostaining and 6-well plates (Falcon) were used to prepare lysates for RNA analyses.

### Virus and infection

TBEV Hypr strain was a kind gift from Dr S. Moutailler (Maisons-Alfort, France). The strain was isolated in 1953 from the blood of a 10-year-old child in the Czech Republic and the complete sequence was published in (71). A working stock was generated in VERO cells (VERO-ATCC-CCL81) cultured in MEM medium (ThermoFisher) supplemented with 2% fetal bovine serum (FBS). Titer was estimated by plaque assay on VERO cells. Neuronal/glial cells differentiated for 13 days were infected with the virus (MOI 10^-2^) for 1h at 37°C. The inoculum was removed and cells were incubated in fresh N2A-NBC medium. Virus titers were estimated by endpoint dilution on VERO cells (TCID50).

### RNA isolation and qPCR

RNA was isolated from infected and non-infected neuronal/glial co-cultures. Cells were lysed using the *NucleoMag® 96 RNA* kit (Macherey Nagel) and RNA was extracted with a *King Fisher Duo* automat (Fisher Scientific) following the manufacturer’s instructions. Extraction of viral RNA from supernatants of infected cells was performed using *QIAamp Viral RNA Mini Kit* (Qiagen) according to the manufacturer’s instructions. One hundred and sixty ng (Fig. 5) or 250 ng (Fig. 7) of RNA were used to synthesize cDNA with the *SuperScript™ II Reverse Transcriptase* kit (ThermoFisher Scientific). Real-time PCR was performed using 2µl of cDNA and *QuantiTect SYBR green PCR master* (Qiagen) with a LightCycler 96 instrument (Roche Applied Science), for a total volume of 20µl of reaction mixture. For relative quantification, the -2ΔΔCt method was used (72). The references genes were *GAPDH* or *HPRT1*. Primers pairs are listed in Supplementary table 1.

### RT² profiler PCR array

Equal volumes of RNA from biological triplicates were pooled for each condition. Two hundred to 500 ng of RNA were transcribed with the *RT² First Strand Kit* (SA Biosciences, Qiagen). Synthetized cDNA was subjected to a PCR array specific for the human antiviral response (*RT² Profiler PCR array* – PAHS-122Z, SA Biosciences, Qiagen), according to the manufacturer’s instructions. Data were normalized using the *HPRT1* house-keeping gene and analyzed with the -2ΔΔCt method for relative quantification. According to the manufacturer’s instructions, an arbitrary cut-off of 3 was applied to determine significant differences. The analysis was performed using the Qiagen Data analysis center (http://www.qiagen.com/fr/shop/genes-and-pathways/data-analysis-center-overview-page/).

### Immunofluorescence assays and cell enumeration

Neuronal/glial cells were fixed for 20 minutes in 4% paraformaldehyde in PBS (Electron Microscopy Sciences) and standard immunofluorescence was performed using antibodies for HuC/HuD (Thermofisher #A21271), βIII-tubulin (Sigma #T8660), GFAP (Dako #M076101-2 or #Z033429-2), OLIG2 (R&D Systems #AF2418) and TBEV-E3. Cells were blocked for 1h in 3% BSA (Sigma), 0.3% Triton-X-100 (VWR) in PBS 1X and primary antibodies were incubated in 1% BSA, 0.1% Triton-X-100 in PBS 1X overnight at +4°C. Secondary antibodies were Alexa Fluor-488/546-conjugated anti-mouse/anti-rabbit IgG (Molecular Probes, Invitrogen). Nuclei were stained with 4′, 6-diamidino-2-phenylindole (DAPI) (Life Technologies) at 0.1 ng/ml. Cell sub-types and infected cells were enumerated either manually or automatically. For manual cell quantification (cells immunostained with antibodies directed against βIII-tubulin and GFAP), images were acquired with an *AxioObserver Z1* (Zeiss) inverted microscope using ZEN software (Zeiss) and analyzed using ImageJ 1.49m software. For automated quantification (cells immunostained with antibodies directed against HuC/HuD, TBEV-E3 and OLIG2 or neurites immunostained with an antibody against βIII-tubulin), images were acquired using the Cellomics ArrayScan automated microscope (Thermofisher Scientific) and analyzed using “Colocalization” or “Neuronal profiling” bio-applications on HCS Studio Cell Analysis Software V6.6.0 (Thermofisher Scientific). In all experiments, an average of 1200 (manual quantification) or 5000 (automated quantification) cells per well were enumerated. The digitized images shown were adjusted for brightness and contrast using ImageJ, without further alteration.

### Magnetic-activated cell sorting

Neuronal/glial cells differentiated for 13 days were detached using Gibco™ TrypLE™ Select Enzyme (1X) and collected into N2A-NBC medium. After centrifugation at 80g for 10 minutes, cells were either sub-cultured (Uns-C) or supplemented with kynurenic acid buffer and sorted according the manufacturer’s instructions using the Microbead Kit (Miltenyi Biotec #130-095-826). In brief, resuspended cells were incubated 10 minutes at 4°C with 20µl biotin-conjugated anti-GLAST (ACSA-1) antibodies per 10^7^ cells, washed and incubated with anti-biotin MicroBeads for 15 minutes at 4°C. Cell sorting was performed using MS columns (Miltenyi Biotec, #130-042-201) placed in a MiniMACS^TM^ separator (Miltenyi Biotec #130-090-312). The cell fractions found in the flow-through or bound to beads were composed of enriched neurons (En-N) and enriched astrocytes (En-As), respectively. Both sorted and unsorted cells were seeded at a density of 100 000 cells per cm² on 24-well µ-plates (IBIDI, #82406) in N2A-NBC or conditioned medium (1:1, fresh N2A-NBC: supernatant of non-infected co-cultures differentiated for 13 days, conditioned for 48h). Conditioned medium allowed neuronal survival in En-N cultures. Half of the medium was replaced every other day.

### Statistical analyses

Data are represented as mean ± standard deviation (SD). Statistical analyses were performed with GraphPad Prism V4.03 or V6.0.1 using an unpaired Student’s t test or a one-way ANOVA analysis (Bonferroni’s Multiple Comparison Test), *=(p<0,05), **=(p<0,01), ***=(p<0,001), non-significant (ns)= (p>0,05).

## Acknowledgments

We are most grateful to Dr S Moutailler for providing the TBEV-Hypr strain. We deeply thank Drs PE Ceccaldi and T Couderc for sharing their insight and Dr N Jouvenet for critical reading of the manuscript.

## Supporting information legends

**S1 Fig.**
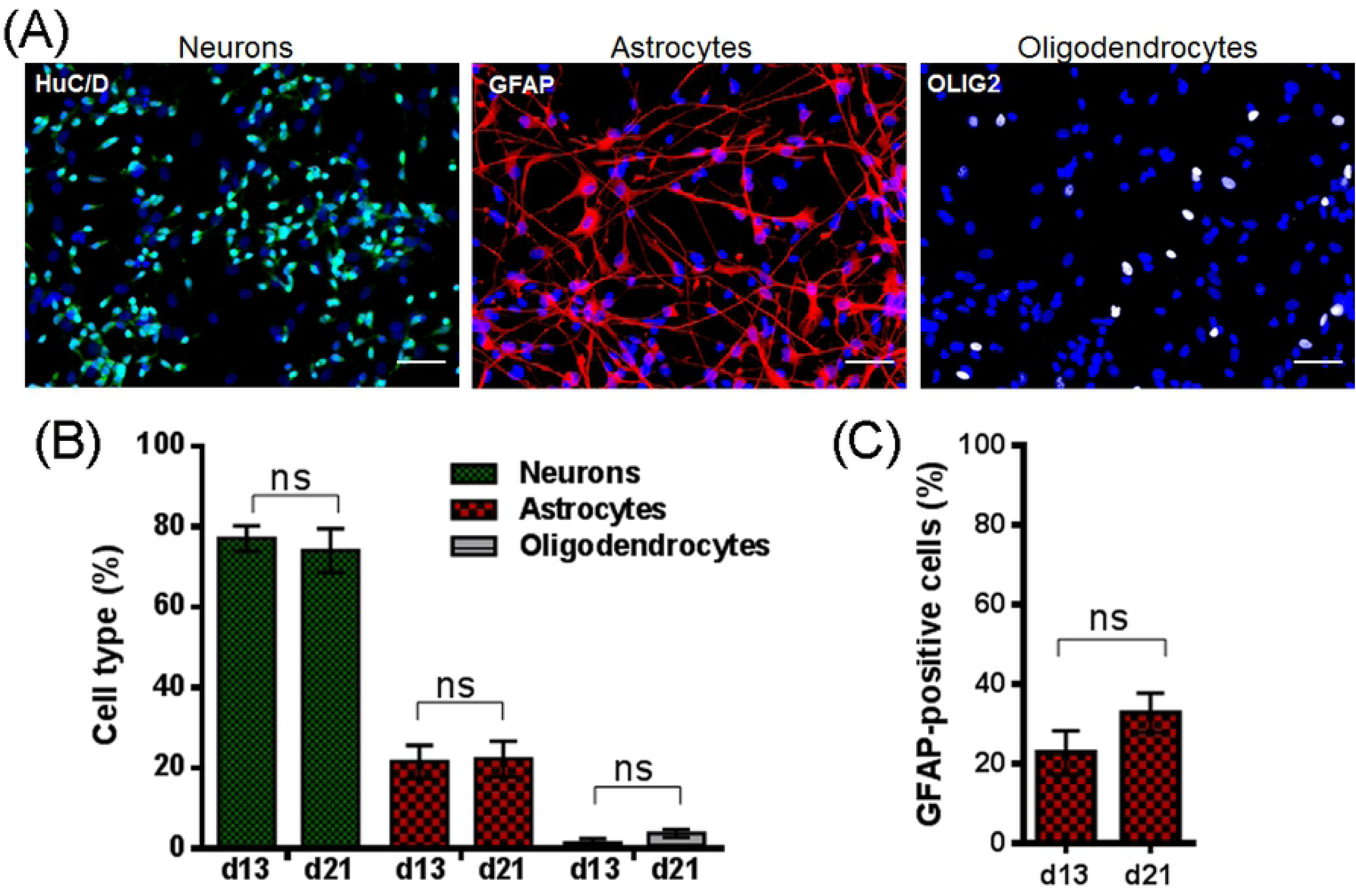
Neurons and astrocytes are the major cell types in hNPCs-derived cultures. HNPCs were differentiated for 13 days. (A) Immunofluorescence labeling using antibodies against HuC/HuD, a neuronal nuclear marker (green), GFAP, an astrocytic marker (red) and OLIG2, an oligodendrocyte nuclear marker (grey) were used. Nuclei were stained with DAPI (blue). Scale bar=20 µm. (B) Enumeration of cells based on immunofluorescence labeling. Automated quantification using an ArrayScan Cellomics instrument. (C) Enumeration of astrocytes based on immunofluorescence labeling. Manual quantification.

**S2 Fig.**
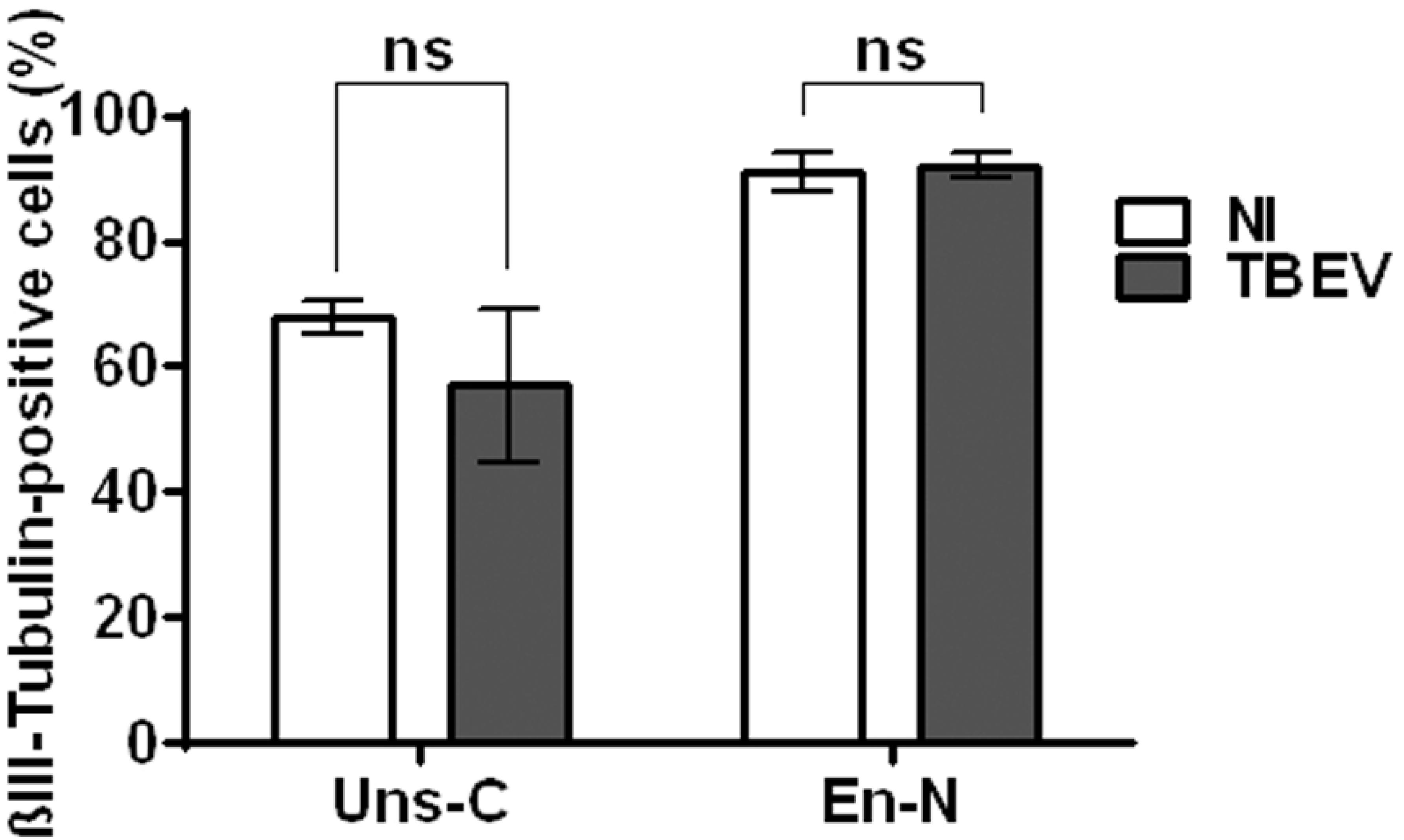
Neuronal survival is not affected by TBEV infection at 24 hpi in unsorted cells and enriched neuron cultures. Unsorted cultures (Uns-C) and enriched neurons (En-N) were infected with TBEV and co-immunostained with βIII-tubulin (neurons) and anti-TBEV-E3 antibodies. Manual enumeration of infected neurons was performed at 24hpi. Data are expressed as the mean±SD and normalized to non-infected Uns-C. Results are representative of two independent experiments performed in triplicate. Statistical analysis was performed using a two-tailed unpaired t test with GraphPad Prism V6.0.1, ns=non-significant (p>0.05).

**Supplementary Table 1.**
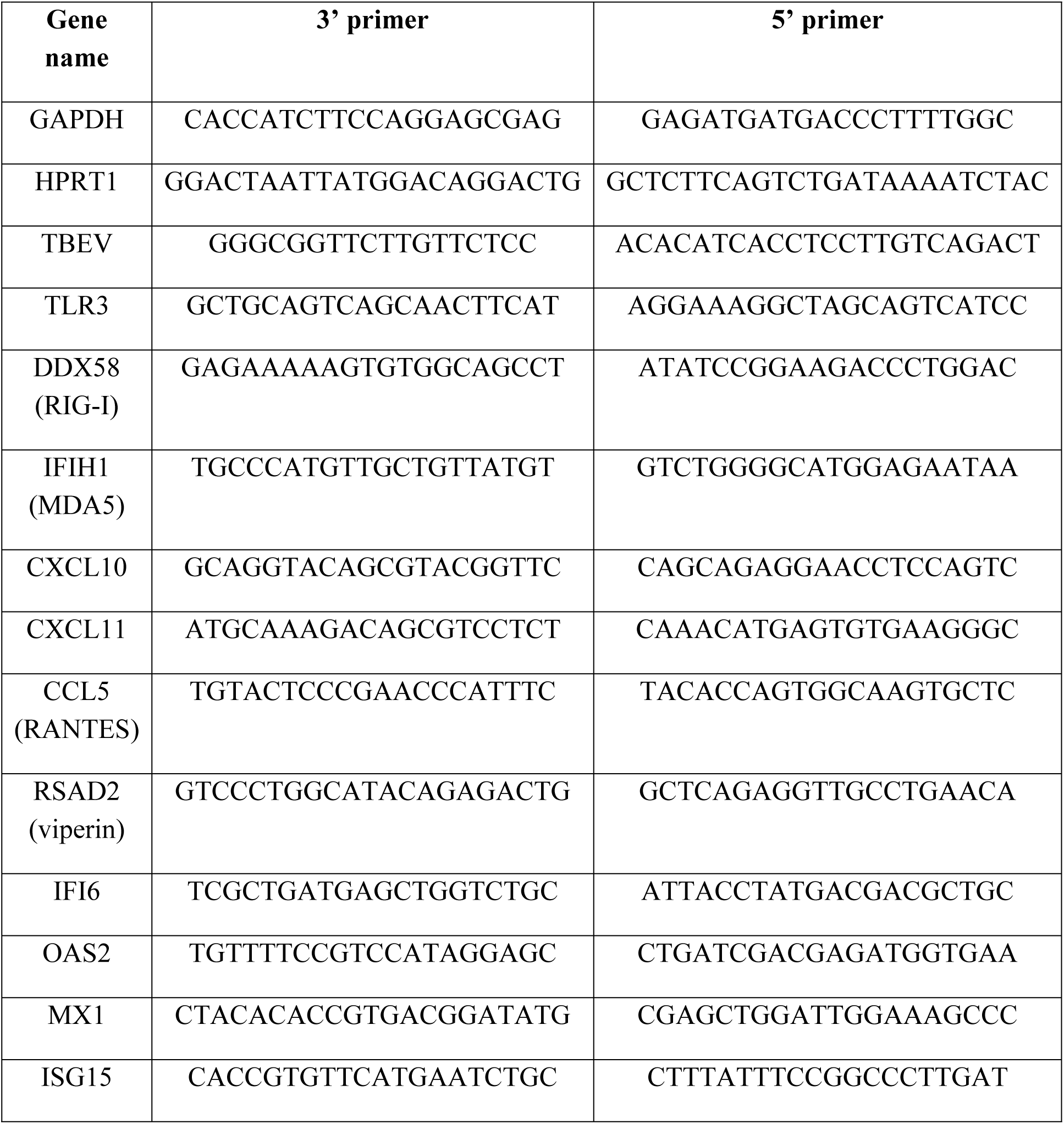
Primer pairs used for qRT-PCR analyses

## Notes

*Funding information:* This study was financially supported by the French National Institute for Agricultural Research (INRA) and DIM MalInf-Ile de France. MF was financially supported by INRA and the Paris Institute of technology for life, food and environmental sciences (AgroParisTech). GG was funded by Labex IBEID. The funders had no role in study design, data collection and analysis, decision to publish, or preparation of the manuscript.

